# MUT-7 exoribonuclease activity and localisation are mediated by an ancient domain

**DOI:** 10.1101/2023.12.20.572533

**Authors:** Virginia Busetto, Lizaveta Pshanichnaya, Raffael Lichtenberger, Stephan Hann, René F. Ketting, Sebastian Falk

## Abstract

The MUT-7 family of 3’-5’ exoribonucleases is evolutionarily conserved across the animal kingdom and plays essential roles in small RNA production in the germline. Most MUT-7 homologs carry a C-terminal domain of unknown function named MUT7-C appended to the exoribonuclease domain. Our analysis shows that the MUT7-C is evolutionary ancient, as a minimal version of the domain exists as an individual protein in prokaryotes. In animals, MUT7-C has acquired an insertion that diverged during evolution, expanding its functions. *C. elegans* MUT-7 contains a specific insertion within MUT7-C, which allows binding to MUT-8 and, consequently, MUT-7 recruitment to germ granules. In addition, in *C. elegans* and human MUT-7, the MUT7-C domain contributes to RNA binding and is thereby crucial for nuclease activity. This RNA-binding function most likely represents the ancestral function of the MUT7-C domain. Overall, this study sheds light on MUT7-C and assigns two functions to this previously uncharacterised domain.

## Introduction

Animals employ RNA interference (RNAi) pathways as a defence mechanism against transposable elements (TEs) and other non-self elements in the germline ^1,2^. This ensures genome integrity and allows the faithful transmission of genetic information to future generations. RNAi is centred around Argonaute (AGO) proteins binding small RNAs (sRNAs) to form the RNA-induced silencing complex (RISC), which recognises by sequence complementarity the target RNA of the non-self elements ^3^. The recognition of the target RNA can trigger transcriptional or post-transcriptional silencing pathways, resulting in gene silencing ^1,2^. To generate an effective RNAi response, animals use different amplification strategies to increase the number of effector sRNAs, also called ‘secondary’ sRNAs, from an initially small pool of ‘primary’ sRNAs that acted as triggers.

In the germline of insects and mammals, a specific class of sRNAs named piRNA are the effectors of RNAi ^4,5^. piRNAs bind to AGO proteins from the PIWI-clade and are amplified through the ping-pong cycle ^6^, which occurs at perinuclear germ granules known as Nuage. Recognition of the TE RNA by a primary RISC results in TE RNA target cleavage. The cleavage product is loaded into a different PIWI protein and subsequently trimmed by the conserved 3’-5’ exoribonuclease Nibbler to become a mature secondary piRNA. The new RISC can then recognise and cleave piRNA precursor transcripts, which are then processed by Nibbler to mature piRNAs ^7–9^.

Nematodes like *C. elegans* employ a different strategy to amplify sRNAs, centred on RNA-dependent RNA polymerases (RdRPs) ^10^. In the *C. elegans* germline, the RdRP RRF-1 localises at perinuclear germ granules named Mutator foci, which are thought to be the site of sRNA amplification ^11–13^. The formation of Mutator foci depends on the intrinsically disordered scaffold protein MUT-16, which acts as a hub for the recruitment of RRF-1 and other proteins to form the Mutator complex ^11,13^. Currently, more than twelve factors have been associated with the Mutator complex and contribute to sRNA amplification. According to the current model, recognition of the target mRNA by the primary RISC results in target cleavage by the endoribonuclease RDE-8 ^14^. Subsequently, the nucleotidyltransferase MUT-2 (also called RDE-3) adds an untemplated polyUG tail to the 3’ end of the newly generated 5’ fragment, marking it for sRNA amplification ^14–16^. This pUG tail is thought to act as a signal for the recruitment of the RdRP RRF-1 to the mRNA fragment that is then used as a template to synthesise a pool of secondary 22G sRNA ^17–19^. These 22G sRNAs are 22 nt long and carry a characteristic guanosine-triphosphate at the 5’ end ^19,20^. Mutations in any Mutator component reduce the level of 22Gs and result in RNAi-resistance and transposon activation in the germline ^11,19,21–23^. While the above model assigns functions to some of the factors of the Mutator complex, the role of most components remains unknown. One of the Mutator factors of yet unknown function is the 3’-5’ exoribonuclease MUT-7, the homolog of *D. melanogaster* Nibbler ^22^. MUT-7 has been proposed to be recruited to MUT-16 by the nematode-specific factor MUT-8 (also called RDE-2) based on *in vivo* studies ^11,24^. MUT-8 has no previously annotated domains, and the mechanism by which it recruits MUT-7 to Mutator foci remains unclear.

Although MUT-7 is highly conserved from sponges to mammals ^8^, a function has only been attributed to Drosophila Nibbler, which trims the 3’ end of pre-piRNAs as part of the ping-pong cycle, but also of subpopulations of miRNAs ^8,9,25,26^. Nibbler is a two-domain protein with an N-terminal domain (NTD) followed by an exoribonuclease (EXO) domain. The NTD consists of a scaffold of HEAT-like repeats with basic patches important for RNA binding and has been proposed to be essential for recruiting Nibbler to AGO-bound RNA substrates ^27^. The Nibbler EXO domain belongs to the DEDDy family of 3′-to-5′ exoribonucleases (InterPro: IPR002562) and is specific to single-stranded RNA ^27^.

In addition to these NTD and EXO domains, *C. elegans* MUT-7 carries an additional C-terminal domain (CTD), absent in Nibbler. This domain is classified as MUT7-C (InterPro: IPR002782) and is predicted to be a deteriorated version of the PIN (PilT N-terminal domain)-like domain fold with an inserted zinc finger at the C-terminus ^28,29^. MUT7-C occurs as a standalone protein in archaea, while bacteria contain two versions: a standalone MUT7-C like in archaea or a fusion to a ubiquitin-like domain ^28,29^. However, to date, no structure of a MUT7-C has been determined, and no function has been assigned.

Here, we show that most Nibbler/MUT-7 homologs carry a MUT7-C domain and that this domain was lost specifically in Drosophilids. We demonstrate that MUT7-C contributes to RNA binding in MUT-7 and its human homolog EXD3. Consequently, deletion of this domain slows down RNA degradation. MUT-7 carries a worm-specific insertion in MUT7-C, which acts as a binding platform for the nematode-specific factor MUT-8. MUT-8, in turn, directly contacts the Mutator scaffold MUT-16, recruiting MUT-7 to Mutator foci. Mutations disrupting the MUT-7/MUT-8 interaction lead to RNAi resistance. These findings assign two functions to the previously uncharacterised MUT7-C domain: an RNA binding function, which most probably represents the most ancient and conserved function and a protein binding platform function that mediates localisation.

## Results

### MUT-7 interacts with the Mutator factor MUT-8

MUT-7 has been reported to interact with the nematode-specific factor MUT-8 by co-immunoprecipitation and yeast two-hybrid experiments ^24^. To understand the mode of interaction between MUT-7 and MUT-8, we analysed the AlphaFold prediction of the two proteins. The exoribonuclease MUT-7 shows an extended, folded N-terminal domain (NTD) followed by an exoribonuclease (EXO) domain and a C-terminal domain (CTD), which is connected to the EXO domain by a flexible linker (**Fig. 1a** and **1b**). MUT-8 is predicted to be mostly unstructured, except for a partially structured NTD that is connected to a structured CTD by a long, flexible linker (**Fig. 1a** and **1b**). While the folded domains of MUT-7 are all predicted with high confidence, all regions of MUT-8 are predicted with low confidence (**Fig. 1b**), potentially due to its restriction to the Caenorhabditis genus (**Supplementary** Fig. 1). Previous yeast two-hybrid experiments showed that MUT-7 residues 643-910 are sufficient to interact with MUT-8 residues 286-578 ^24^. These minimal constructs roughly correspond to the MUT-7 CTD and MUT-8 CTD. To test if MUT-7 directly binds MUT-8 and if the two CTDs mediate the interaction, we performed co-expression pulldown assays with the following constructs MUT-7^FL^ (1-910), MUT-7^NTD-EXO^ (1-625), MUT-7^CTD^ (633-910), MUT-8^FL^ (1-578) and MUT-8^CTD^ (322-578). MUT-7 constructs contained an N-terminal MBP-tag and served as bait. MUT-8^FL^ co-precipitated with MUT-7^FL^ but not with MUT-7^NTD-EXO^, and both MUT-8^FL^ and MUT-8^CTD^ also co-precipitated with MUT-7^CTD^ (**Fig. 1c**). This suggests that the two full-length proteins interact directly and that MUT-7^CTD^ and MUT-8^CTD^ mediate the interaction. We note that MUT-8^FL^ and MUT-8^CTD^ are insoluble without MUT-7, suggesting MUT-8’s solubility depends on MUT-7 (**Fig. 1c**). To corroborate those findings, we purified the MUT-7^FL^/MUT-8^FL^ complex through several chromatographic steps, showing that MUT-7^FL^ and MUT-8^FL^ co-migrate in size exclusion chromatography (SEC) (**Fig. 1d**). Analysis of the purified minimal MUT-7^CTD^/MUT-8^CTD^ complex by SEC-Multi-Angle Light Scattering showed a single peak and an average molecular mass of 60 kDa, confirming that MUT-7^CTD^ and MUT-8^CTD^ form a heterodimer (**Supplementary** Fig. 2a).

**Figure 1:**
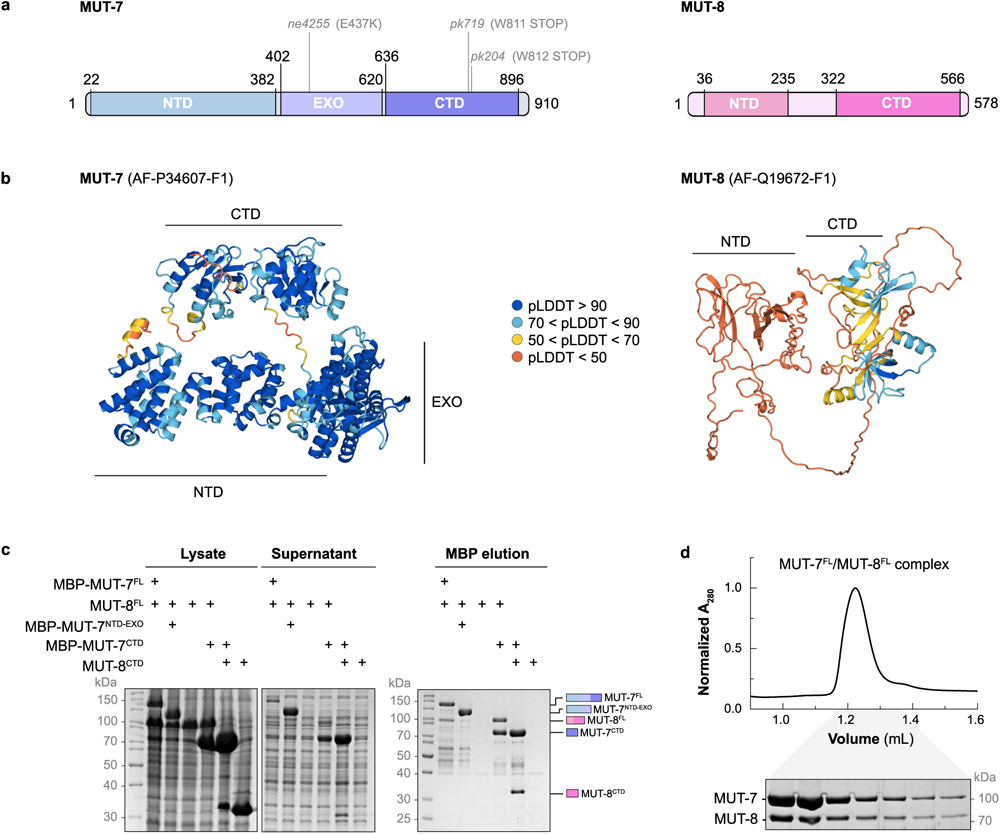
MUT-7 forms a complex with the Mutator factor MUT-8. **a**, Schematic drawing of MUT-7 (P34607) and MUT-8 (Q19672) proteins. Coloured rectangles highlight folded domains. *mut-7* alleles mentioned in this study are indicated. **b**, AlphaFold (AF) prediction of MUT-7 (AF-P34607-F1) and MUT-8 (AF-Q19672-F1). Proteins are coloured according to the pLDDT score of their AlphaFold prediction. A high score indicates high confidence of the AF model, while a low score corresponds to low model confidence. **c**, Analysis of the interaction between MUT-7 and MUT-8 constructs by MBP-pulldown assays. MBP-tagged MUT-7^FL^, MUT-7^NTD-EXO^ and MUT-7^CTD^ were co-expressed with MUT-8^FL^ or MUT-8^CTD^ in *E. coli*. Expression of the preys alone (MUT-8^FL^ and MUT-8^CTD^) is used as a negative control. Total lysate, supernatant and elution are analysed by SDS-PAGE, followed by Coomassie staining. **d**, Size exclusion chromatography analysis of the MUT-7^FL^/MUT-8^FL^ complex on a Superdex 200 increase 3.2/300 column. The peak fractions are visualised on a Coomassie-stained SDS polyacrylamide gel.

### Crystal structure of the MUT-7^CTD^/MUT-8^CTD^ complex

Initial attempts to crystallise the MUT-7^CTD^/MUT-8^CTD^ heterodimer were unsuccessful. Limited proteolysis with different proteases resulted in a similar degradation pattern, with MUT-8^CTD^ left intact and MUT-7^CTD^ cleaved into two shorter fragments (**Supplementary** Fig. 2b and 2c). MUT-8^CTD^ and the two MUT-7^CTD^ fragments co-migrate during SEC, indicating that the complex is stable (**Supplementary** Fig. 2d). Further trimming of a few external amino acids on CTDs of MUT-7 and MUT-8 yielded diffracting crystals with one copy of the complex in the asymmetric unit. The phases were determined by molecular replacement using the MUT-7^CTD^ AlphaFold model and refined with an R-free of 22.1%, R-factor of 18.8%, and good stereochemistry (**Table S1,** PDB ID: 8Q66).

MUT-7^CTD^ and MUT-8^CTD^ are elongated molecules interacting via an extended interface of 2140 Å^2^ (Fig. 2a). The MUT-7^CTD^ can be subdivided into an N-terminal domain (MUT-7^CTD-N^ (633-767)) and a C-terminal domain (MUT-7^CTD-C^ (768-899)) containing a zinc finger with four cysteines coordinating a Zn^2+^ ion (Fig. 2b and 2c). We used Foldseek ^30^ and DALI ^31^ to identify domains with structural similarity. MUT-7^CTD-N^ is similar to the response regulator receiver protein from *B. phymatum* ^32^ and the TOPRIM domain of RNase M5 from *G. stearothermophilus* ^33^ (**Supplementary** Fig. 3a-c). For MUT-7^CTD-C^, we could not find structures with a Template modelling (TM)-score higher than 0.4, suggesting that this fold is not yet represented by experimentally determined structures in the PDB (**Supplementary** Fig. 3a). Previous bioinformatic analysis based on sequence comparison had described MUT-7 CTD as a deteriorated version of the PIN-like domain with an additional zinc finger fused at the C-terminus ^28^. Structural comparison of MUT-7^CTD-N^ against the PDB with Foldseek ^30^ and DALI ^31^ did not identify any structurally related PIN domains. Manual comparison of the MUT-7^CTD-N^ with two representative PIN domains from Nob1 and VapC (PDB: 6TG6 and 4CHG) using TM-align ^34^ resulted in TM-scores of around 0.3, indicating that they are only distantly related.

**Figure 2:**
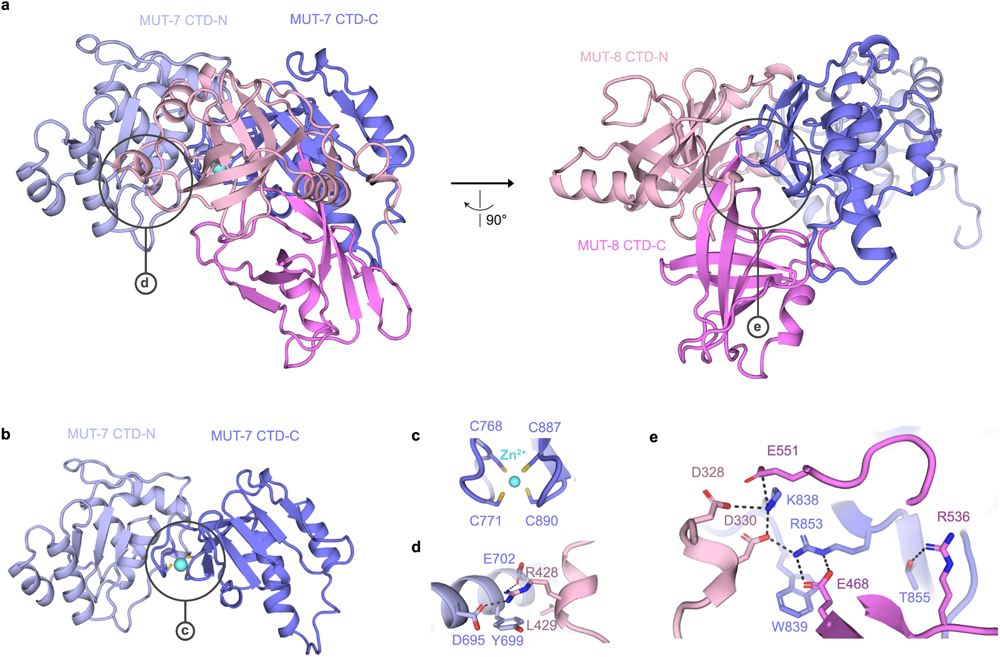
Crystal structure of the MUT-7^CTD^/MUT-8^CTD^ complex. Cartoon representation of the MUT-7^CTD^/MUT-8^CTD^ complex (PDB ID: 8Q66). The respective CTD-N and CTD-C subdomains are shown in shades of blue (MUT-7^CTD^) or pink (MUT-8^CTD^). The Zn^2+^ ion is in cyan. **a**, Two different orientations of the complex. Black circles highlight the main interaction surfaces between MUT-7^CTD^ and MUT-8^CTD^, of which a close-up view is shown in (d) and (e). **b,** Cartoon visualisation of MUT-7^CTD^. A black circle highlights the Cys4 zinc finger, of which a close-up view is shown in (c). **c**, Close-up view of MUT-7^CTD-C^ Cys4 zinc finger. **d**, Close-up view of the interaction between MUT-7^CTD-N^ and MUT-8^CTD-N^. **e**, Close-up view of the interaction between MUT-7^CTD-C^ and MUT-8^CTD^.

The MUT-8^CTD^ can also be divided into two subdomains, MUT-8^CTD-N^ and MUT-8^CTD-C^ (Fig. 2a and **Supplementary** Fig.1). Structural analysis using Foldseek ^30^ revealed that both subdomains show similarity to oligonucleotide/oligosaccharide-binding (OB)-folds flanked by additional secondary structure elements (**Supplementary** Fig. 3a and 3d**-g**). OB-fold domains can mediate protein-RNA as well as protein-protein interactions (PMID: 12504685).

Both the MUT-7 and MUT-8 CTD-N and CTD-C subdomains contribute to complex formation. On one side, a short helical insertion in MUT-8^CTD-N^ (Arg428, Leu429) interacts with MUT-7^CTD-N^ (Asp695, Tyr699 and Glu702) (Fig. 2a and 2d). This interface is supported by crosslinking mass spectrometry of the MUT-7^CTD^/MUT-8^CTD^ complex, which shows crosslinks between K701 of MUT-7 and T425 or K426 of MUT-8 (**Supplementary** Fig. 2e). On the other side, MUT-7^CTD-C^ forms an extended interaction network with both MUT-8^CTD-N^ and MUT-8^CTD-C^. MUT-7 Arg853 interacts electrostatically with MUT-8 Glu468 and Asp330. MUT-8 Asp330, in turn, interacts with MUT-7 Lys838, which is further stabilised by the interaction with MUT-8 Asp328 and Glu551. Next to MUT-7 Arg853, Thr855 forms a hydrogen bond to MUT-8 Arg536 (Fig. 2a and 2e). Overall, MUT-7^CTD-N^ and MUT-7^CTD-C^ contribute to MUT-8 binding, with MUT-7^CTD-C^ playing a major role. We tested if MUT-7^CTD-C^ was sufficient for MUT-8^CTD^ binding by co-expressing MUT-8^CTD^ with either MBP-tagged MUT-7^CTD^ or MUT-7^CTD-C(773–910)^ and analysed the interaction by pulldown assays. The MUT-8^CTD^ co-precipitated poorly with MUT-7^CTD-C(773–9^^10)^, indicating that MUT-7^CTD-C^ is necessary but insufficient for binding MUT-8 (**Supplementary** Fig. 3h).

Overall, these results give structural insights into a MUT7-C domain and assign a protein-binding function to this previously uncharacterised domain.

### MUT-8 bridges MUT-7 to the Mutator scaffold protein MUT-16

MUT-7 and MUT-8 co-immunoprecipitate and co-localize with the Mutator scaffold factor MUT-16 and are therefore considered part of the Mutator complex ^11,13,35^. Based on the observation that MUT-7 fails to localise to germ granules in a *mut-8(pk1657)* mutant strain ^11^, MUT-8 has been proposed to recruit MUT-7 to Mutator foci. However, no direct interaction between MUT-16 and MUT-8 or any other Mutator component has yet been shown. Of note, two MUT-16 isoforms are annotated: B0379.3a.1 (Uniprot: O62011) and B0379.3b.1 (Uniprot: Q9U3S5), the latter lacking four amino acids (122-Arg-Asp-Leu-Gln-125). Here, we use the longer B0379.3a.1 isoform as a reference, and adapted nomenclature accordingly when referring to experiments using B0379.3b.1 as reference ^11,13^. The AlphaFold prediction shows that MUT-16 is mostly disordered with only two structured domains, one at the N-terminus and a small one at the C-terminus. A region comprising the C-terminal part of the IDR together with the folded C-terminal domain (777-1037) is necessary and sufficient for foci formation in worms ^13^ (Fig. 3a).

**Figure 3:**
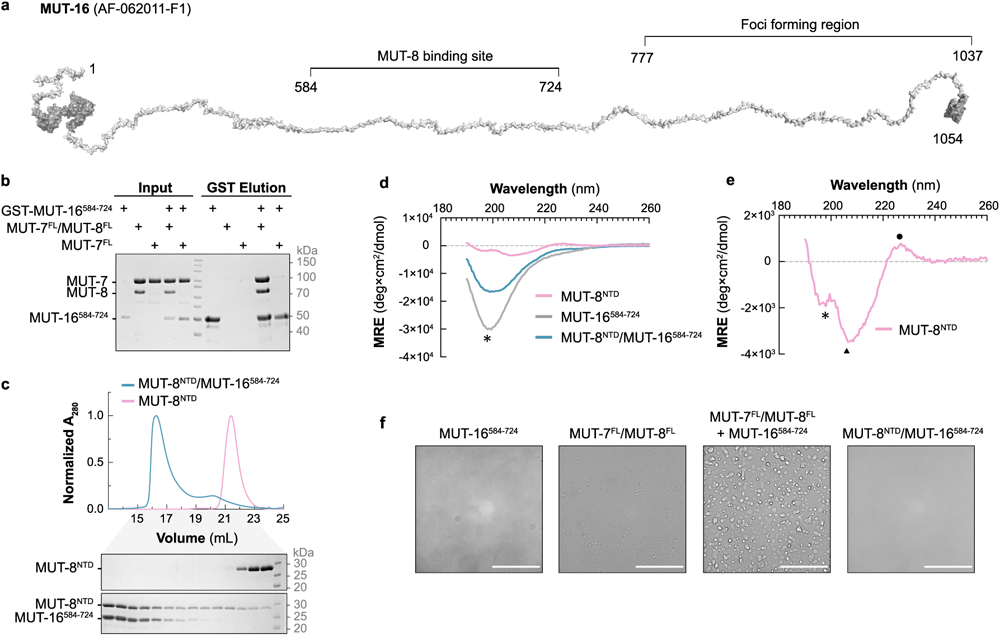
The ternary complex MUT-7/MUT-8/MUT-16 forms condensates. **a**, MUT-16 AlphaFold prediction shown as cartoon (AF-O62011-F1), stretched with PyMOL. Folded regions are in dark grey and disordered regions in light grey. The MUT-8 binding region identified in this study (584-724) and the foci-forming region (777-1037) are highlighted. **b**, GST pulldown assay with purified proteins to test the interaction between GST-tagged MUT-16^584–724^ and MUT-7^FL^ or the MUT-7^FL^/MUT-8^FL^ complex. Proteins were incubated with glutathione-coupled beads. Input and elution fractions were analysed by SDS-PAGE followed by Coomassie staining. **c**, Size exclusion chromatography analysis of purified MUT-8^NTD^ and the complex MUT-8^NTD^/MUT-16^584–724^) on a Superdex 200 increase 10/300 GL column (Cytiva). Chromatograms: MUT-8^NTD^ (pink), MUT-8^NTD^/MUT-16^584–724^ (petrol green). The Coomassie-stained SDS polyacrylamide gels show the peak fractions from size exclusion chromatography. **d** and **e**, Far-UV CD spectra of MUT-8^NTD^ (pink), MUT-16^584–724^ (grey) and the MUT-8^NTD^/MUT-16^584–724^complex (petrol green). MRE = Mean Residue Ellipticity. A zoom-in on the CD spectrum of MUT-8^NTD^ is shown in (e). Peaks with a minimum at 197/198 nm, indicative of an intrinsically disordered protein, are marked by an asterisk. The weak positive peak at 226 nm and a pronounced minimum near 207 nm, indicative of polyproline II conformation, are marked by a circle and a triangle, respectively. **f**, Phase separation assays using bright-field imaging at a protein concentration of 10 µM in the presence of 5% (w/v) PEG6000 as a crowding agent. The proteins used are indicated above the image. Scale bars correspond to 50 μm.

To test if MUT-16 directly contacts MUT-7/MUT-8 and to determine the interacting regions, we performed pulldown experiments. As we failed to express full-length MUT-16, we designed three C-terminally truncated MUT-16 constructs (1-724, 1-584 and 1-383) based on multiple sequence alignment (**Supplementary** Fig. 4) and purified them as GST-tagged fusion proteins for pulldown experiments with full-length MUT-7/MUT-8. While the longest MUT-16 construct (MUT-16^1–7^^24^) co-precipitated MUT-7/MUT-8, the two shorter MUT-16 constructs (MUT-16^1–584^ and MUT-16^1–383^) failed to bind MUT-7/MUT-8 (**Supplementary** Fig. 5a), suggesting that the MUT-16 region 584-724 is essential for MUT7/MUT-8 binding. We purified MUT-16^584–724^ and tested its binding to MUT-7^FL^/MUT-8^FL^ in pulldown experiments. MUT-7^FL^/MUT-8^FL^ co-precipitated with GST-tagged MUT-16^584–724^ as efficiently as with MUT-16^1–724^, indicating that this MUT-16 region is necessary and sufficient for the interaction (Fig 3b and **Supplementary** Fig. 5b). In contrast to full-length MUT-7/MUT-8, the MUT-7^CTD^/MUT-8^CTD^ complex did not bind MUT-16^584–724^, indicating that MUT-16 binding is not mediated by the CTD of either MUT-7 or MUT-8 (**Supplementary** Fig. 5b).

To investigate if MUT-7/MUT-8 interaction with MUT-16 depends on MUT-8, we purified MUT-7^FL^ and tested its binding to GST-MUT-16^584-724^. Without MUT-8, MUT-7^FL^ did not co-precipitate with MUT-16^584-724^ (Fig. 3b). Our results indicate that MUT-8 is necessary for MUT-16 binding, but MUT-8 C-terminal domain is insufficient, suggesting that upstream regions might mediate MUT-16 binding. According to AlphaFold, MUT-8 has a partially structured NTD (36-235) (Fig. 1b), which we hypothesised interacts with MUT-16. Co-expression of MUT-16^584-724^ with MBP-tagged MUT-8^FL^, MUT-8^NTD^ (1-235) or MUT-8^CTD^ followed by MBP pulldown showed that MUT-16^584-724^ directly binds to the NTD of MUT-8 (**Supplementary** Fig. 5c). Moreover, analysis of the MUT-8^NTD^/MUT-16^584-724^complex by SEC confirmed the direct interaction of the two constructs (Fig. 3c). Quantitative analysis of the interaction between MUT-8^NTD^ and MUT-16^584-724^ using isothermal titration calorimetry revealed a *K*_d_ around 13 μM (**Supplementary** Fig. 5d).

Our findings indicate that MUT-8 uses its folded CTD to bind the MUT-7 CTD and its NTD to contact MUT-16, thus bridging MUT-7 and MUT-16. In agreement with our results, a previous study showed that MUT-8 fails to localise correctly at Mutator foci in worms carrying a MUT-16 deletion in a region (636-776) comparable to the one we identified as MUT-8 binding site ^13^.

### MUT-7 recruitment to germ granules involves condensate formation

AlphaFold predicts the region of MUT-16 that binds MUT-8 to be fully disordered (Fig. 3a), while the MUT-8^NTD^ is predicted to be partially structured, containing some β-strands (Fig. 1b).

We purified untagged MUT-16^584-724^ and used circular dichroism (CD) spectroscopy to analyse the secondary structure content of MUT-16^584-724^, MUT-8^NTD^ and the MUT-8^NTD^/MUT-16^584-724^ complex. The far-UV CD spectrum of MUT-16^584–724^ shows a minimum at 197 nm, indicative of an intrinsically disordered region (Fig. 3d). The spectrum of MUT-8^NTD^ shows a weak positive peak at 226 nm, a pronounced minimum near 207 nm, and a shoulder at 197 nm (Fig. 3e). The peaks at 226 and 207 nm are reminiscent of polyproline II conformation ^3^^6^, and the shoulder at 197 nm of a disordered polypeptide, suggesting that MUT-8^NTD^ is a mixture of a disordered polypeptide with the presence of regions in a polyproline II conformation. The spectrum of the purified MUT-8^NTD^/MUT-16^584–724^ complex shows a broader minimum at around 200 nm, suggesting that the complex has low secondary structure content. Thus, complex formation seems not to be coupled to secondary structure formation, as the spectrum of the complex can be explained by the sum of the individual spectra (Fig. 3d).

The MUT-16 C-terminal region (777-1037) is necessary and sufficient for foci formation *in vivo*, but N-terminally extended MUT-16 constructs result in an increased number and size of Mutator foci ^1^^3^. We thus investigated the propensity of MUT-16^584–724^ to form condensates in the presence or absence of MUT-7/MUT-8. While MUT-16^584–724^ and MUT-7^FL^/MUT-8^FL^ on their own did not form condensates, the ternary MUT-7^FL^/MUT-8^FL^/MUT-16^584–724^ complex formed droplets at a concentration of 10 µM in the presence of 5% (w/v) PEG6000 as a crowding agent. Interestingly, the minimal complex MUT-8^NTD^/MUT-16^584–724^ did not form droplets, suggesting additional contributions from the MUT-8^CTD^ or MUT-7 or both to condensate formation (Fig. 3f). We conclude that MUT-7/MUT-8 binding to MUT-16 promotes condensation and thereby contributes to Mutator foci formation.

### The MUT-7/MUT-8 interaction is required for functional RNA interference

Our biochemical data suggest that MUT-8 acts as an adapter, bridging MUT-7 to MUT-16 via condensate formation (Fig. 4a). To confirm that MUT-8 recruits MUT-7 to germ granules, we designed mutations at the MUT-7^CTD^/MUT-8^CTD^ interface that would disrupt MUT-7/MUT-8 interaction. When MUT-7^CTD^ Arg853 and Thr855 were mutated to glutamate residues, MUT-8^CTD^ failed to co-precipitate with MBP-tagged MUT-7^CTD^, indicating that the selected mutations disrupt the interaction (Fig. 4b). We introduced these two MUT-7 mutations (R853E, T855E) in worms and tested the resulting strain for RNAi proficiency. As a positive control we used a worm strain (*pk204*, see Figure 1a) carrying a nonsense mutation in the *mut-7* gene, which was previously shown to be RNAi resistant ^2^^1,2^^2^. We found that the selected mutations (R853E, T855E) result in fully RNAi-resistant animals (Fig. 4c). Mutator foci are still formed in the *mut-7* (R853E, T855E) mutant animals, indicating that the observed phenotype is not due to Mutator foci disruption (Fig. 4d). We conclude that the identified interface between MUT-7 and MUT-8 is important for MUT-7 function *in vivo*.

**Figure 4:**
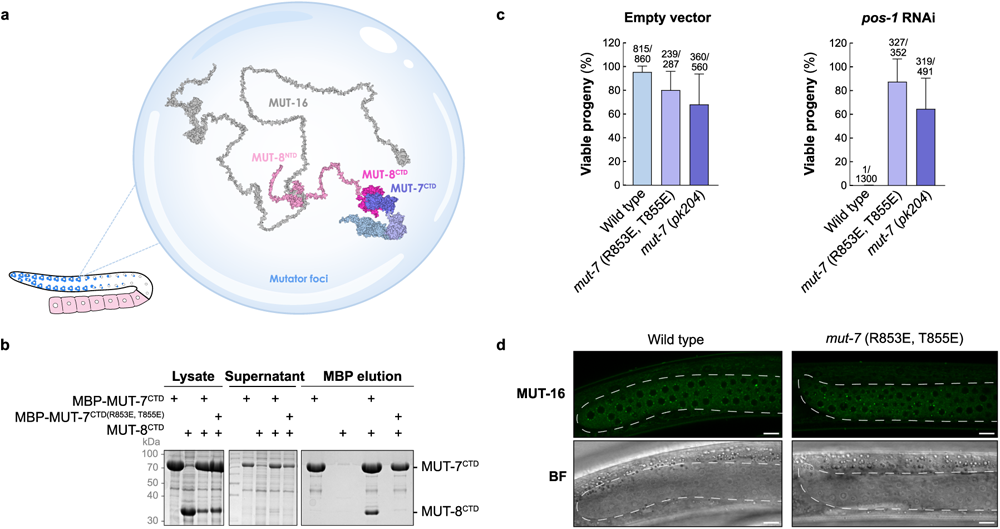
MUT-7 binding to MUT-8 is required for functional RNAi. **a,** Schematic of MUT-7 recruitment to Mutator foci by MUT-8. The cartoon showing *C. elegans* germline was adapted from Aoki et al. ^3^^7^. A zoom-in illustrates one Mutator focus. MUT-16, MUT-8 and MUT-7 cartoon representations correspond to the isolated AlphaFold predictions. The interaction between proteins has been modelled with PyMOL based on our biochemical results and it is not a prediction. **b**, Pulldown assay to test the impact of MUT-7^CTD^ (R853E, T855E) mutations on MUT-8^CTD^ binding. MBP-tagged MUT-7^CTD^ or MUT-7^CTD(R853E, T855E)^ were co-expressed with MUT-8^CTD^ in *E. coli*. Expression of MUT-8^CTD^ alone served as negative control. After lysis, the supernatant is incubated with amylose resin. Total lysate, supernatant and MBP-elution were analysed by SDS-PAGE followed by Coomassie staining. **c**, Percentage of viable progeny for worms fed with bacteria carrying an empty vector or a vector expressing a dsRNA against *pos-1*. Three *C. elegans* strains were tested: a wild-type (WT) strain, a strain carrying the MUT-7^CTD^ (R853E, T855E) mutation, and the previously characterized *mut-7*(*pk204*) strain. The number of hatched larvae/total number of laid eggs used to calculate the percentage of viable progeny for each strain is indicated above the bars. Error bars represent ± SD of the mean. **d**, Expression and localization of MUT-16::GFP::FLAG in wild type and *mut-7* (R853E, T855E) worm strains. The mitotic part of the young adult gonad is shown. A representative image from a total of three analysed animals is shown. BF: Bright Field. Scale bars: 10 μm.

### MUT-7 is a metal-dependent 3’-5’ exoribonuclease

Although the role of MUT-7 in the sRNA amplification process remains unknown, its 3’-5’ exoribonuclease activity is necessary for the production of secondary sRNAs, as a mutation in its catalytic site (E437K) leads to TE mobilisation ^1^^9^. The AlphaFold prediction of MUT-7 reveals that the EXO domain adopts an RNase D-like fold with a deep catalytic pocket containing the four conserved residues (DEDD) (Fig. 5a), like its *D. melanogaster* homolog Nibbler ^27^. To test the exoribonuclease activity of MUT-7, we incubated recombinant MUT-7^FL^ with a 5′-6-fluorescein amidite (5′-FAM)-labelled 28-mer ssRNA. Reaction products were detected by fluorescence on 15% TBE-Urea polyacrylamide gels. MUT-7^FL^ showed the highest 3′-5′ exoribonuclease activity in the presence of Mn^2+^, some activity in the presence of Mg^2+^, and no exonuclease activity with other divalent metal ions such as Zn^2+^ and Ca^2+^ (Fig. 5b), as previously shown for Nibbler ^27^. We then tested the 3’-5’ exoribonuclease activity of the MUT-7^FL^/MUT-8^FL^ complex to see if MUT-8 impacts MUT-7’s activity. Based on the Nibbler D435A mutant ^27^, we engineered and purified a catalytically inactive MUT-7^FL(D453A)^/MUT-8^CTD^ complex as a negative control. The MUT-7^FL^/MUT-8^FL^ complex degraded ssRNA with similar efficiency as MUT-7^FL^ alone, while MUT-7^FL(D453A)^/MUT-8^CTD^ was inactive. We conclude that MUT-8 does not affect MUT-7 exoribonuclease activity (Fig. 5c).

**Figure 5:**
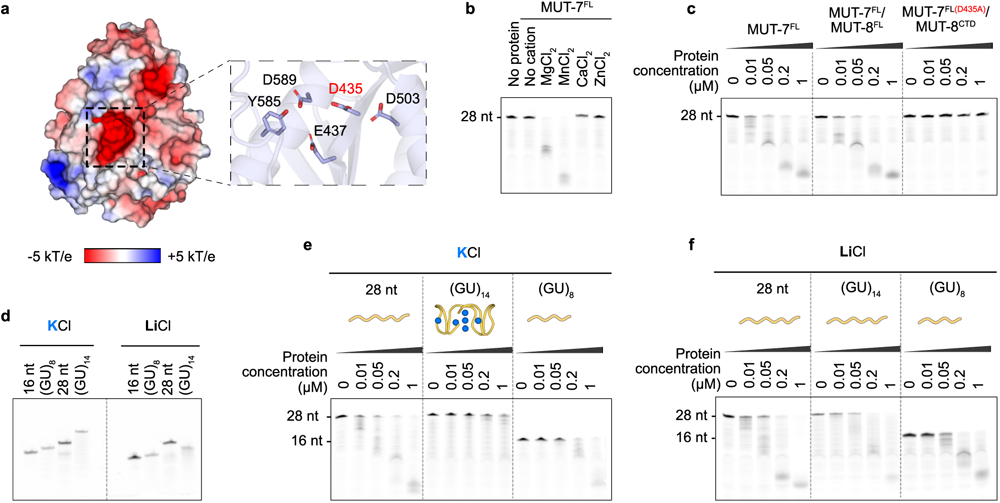
MUT-7 degrades ssRNA but not a pUG fold in a metal-dependent manner. **a**, Electrostatic surface potential of MUT-7 exoribonuclease domain based on the AlphaFold prediction. Electrostatic surface potential was calculated by Adaptive Poisson-Boltzmann Solver (APBS) from −5 kT/e (red) to +5 kT/e (blue). The zoom-in shows the MUT-7 catalytic site; the mutated Asp435 in inactive MUT-7 is highlighted in red. **b**, Metal ion-dependency of MUT-7 3′-5′ exoribonuclease activity. Purified MUT-7 (0.5 μM) was incubated with a 28-mer ssRNA carrying 5′-FAM-labeled 28-mer ssRNA (1 μM) for 30 minutes at room temperature in the absence or presence of different divalent cations (5 mM MgCl_2_, MnCl_2_, ZnCl_2_, or CaCl_2_). Reaction products were separated by TBE-Urea PAGE. **c**, Comparison of MUT-7 and MUT-7/MUT-8 exoribonuclease activity by a dose-dependent nuclease assay. An inactive MUT-7^D435A^/MUT-8^CTD^ complex was used as negative control. Increasing amounts of proteins were incubated with a 5′-FAM-labeled 28-mer ssRNA (1 μM) in the presence of 2 mM MnCl_2_ for 30 minutes at room temperature. Reaction products were separated by TBE-Urea PAGE. **d**, Native PAGE of indicated 5′-FAM-labeled RNAs in the presence of either 100 mM KCl or LiCl. (GU)_8_ and (GU)_14_ RNAs were pre-folded in folding buffer. A 16-mer and a 28-mer RNAs were used as control. **e**-**f**, Dose-dependent exoribonuclease assays with a pre-folded (GU)_14_ RNA in a buffer containing 50 mM KCl (e) or 50 mM LiCl, preventing pUG fold formation (f). A cartoon representation of the linear RNAs or the pUG fold (PDB: 7MKT) is shown. Potassium ions are in blue. Increasing amounts of MUT-7 were incubated with a pre-folded (GU)_14_ RNA, a (GU)_8_ RNA that cannot adopt the pUG fold and a 28-mer ssRNA. All RNAs were 5′-FAM-labeled. Reaction products were separated by TBE-Urea PAGE.

We next investigated MUT-7 substrate preferences to get hints on MUT-7 putative role in sRNA amplification. In Drosophila, Nibbler trims the 3’ end of both miRNAs and piRNAs ^8,25,26^. Similarly, *C. elegans* MUT-7 might be necessary for trimming RdRP products to their final length of 22 nt. Alternatively, MUT-7 could act on different RNA substrates. A prominent RNA structure that plays a critical role in the *C. elegans* sRNA amplification process is the pUG fold ^15,17,18,38^. Following template RNA cleavage, the newly generated 3’ hydroxyl is modified by the nucleotidyltransferase MUT-2, which appends repeating dinucleotide UG-units ^15^. In the presence of potassium ions, 11.5 UG/GU repeats can form a compact quadruplex (G4) structure named pUG fold ^18,39^. We tested MUT-7’s ability to degrade (GU)_8_ and (GU)_14_ RNAs, with the (GU)_14_ RNA being long enough to adopt a pUG fold in the presence of potassium but not lithium ions ^18,39^, as shown by native PAGE (Fig. 5d). As a control, we used a 28-mer ssRNA incapable of adopting a pUG fold. MUT-7^FL^ efficiently degraded the (GU)_8_ and the 28-mer ssRNA but failed to efficiently degrade the (GU)_14_ RNA in the presence of potassium ions (Fig. 5e). To confirm that RNA folding prevents degradation, we repeated the assay with lithium instead of potassium ions. Strikingly, MUT-7^FL^ was able to degrade all RNA substrates in the presence of lithium ions, indicating that folded RNAs cannot be degraded by MUT-7 (Fig. 5f).

### The MUT7-C domain is evolutionarily conserved throughout all domains of life

We showed that *C. elegans* MUT-7_CTD_ binds MUT-8_CTD_, which in turn interacts with MUT-16 through its NTD. Hence, one role of the MUT-7 CTD in *C. elegans* is to establish the MUT-7 localisation to Mutator foci via MUT-8. Given that both MUT-8 and MUT-16 are conserved in Caenorhabditis (Supplementary Fig. 1 and 4), this MUT-7 CTD function is likely to be extended to other Caenorhabditis species. Interestingly, the only MUT-7 homolog that has been characterised is *D. melanogaster* Nibbler, which lacks the domain classified as MUT7-C. A comprehensive analysis of the presence/absence of the MUT7-C domain in MUT-7 animal homologs has not yet been done. We performed a thorough phylogenetic analysis of the distribution of MUT-7 proteins in animals. This revealed that most animal homologs, including the vertebrate factor EXD3, carry a MUT7-C domain downstream of the EXO domain. The MUT7-C domain is also present in the common ancestors of Nibbler family members, indicating that a loss has occurred in Drosophilids (**Supplementary** Fig. 6). Interestingly, in the yellow fever mosquito (*A. aegypti*) Nibbler does carry the MUT7-C (A0A7N4YH63). However, recent work did not consider this domain ^27^, likely because the previously annotated isoform lacked the CTD (Q179T2). Since Mutator complexes are not present in animals besides Caenorhabditis ^11,13^, this implies that the MUT7-C domain likely has a function beyond mediating MUT-8 binding. The MUT7-C domain is evolutionarily ancient, being also found as an individual protein in archaea and bacteria. In some bacteria, it is combined with an N-terminal ubiquitin-like domain ^28,29^. So far, no function has been assigned to MUT7-C domains. To identify the core elements of the MUT7-C fold and gain insights into the potentially conserved function of the MUT7-C domain, we generated a multiple sequence alignment including bacterial, archaeal, and eukaryotic sequences (Fig. 6a **and** **Supplementary** Fig. 7a). In parallel, we compared the experimental structure of *C. elegans* MUT-7 CTD with the AlphaFold predictions of CTDs from human EXD3, *A. aegypti* Nibbler and prokaryotic MUT7-C domains (Fig. 6b). This revealed that prokaryota and plants carry a minimal conserved MUT7-C unit consisting of an N-terminal part (CTD-N) and a C-terminal part (CTD-C). We used the Alphafold structure of the Archaeon *Thermococcus eurythermalis* as representative of the most minimal MUT7-C fold to visualise the conservation level of the different elements. A detailed analysis of the core structural elements is shown in **Supplementary** Fig. 7b.

**Figure 6:**
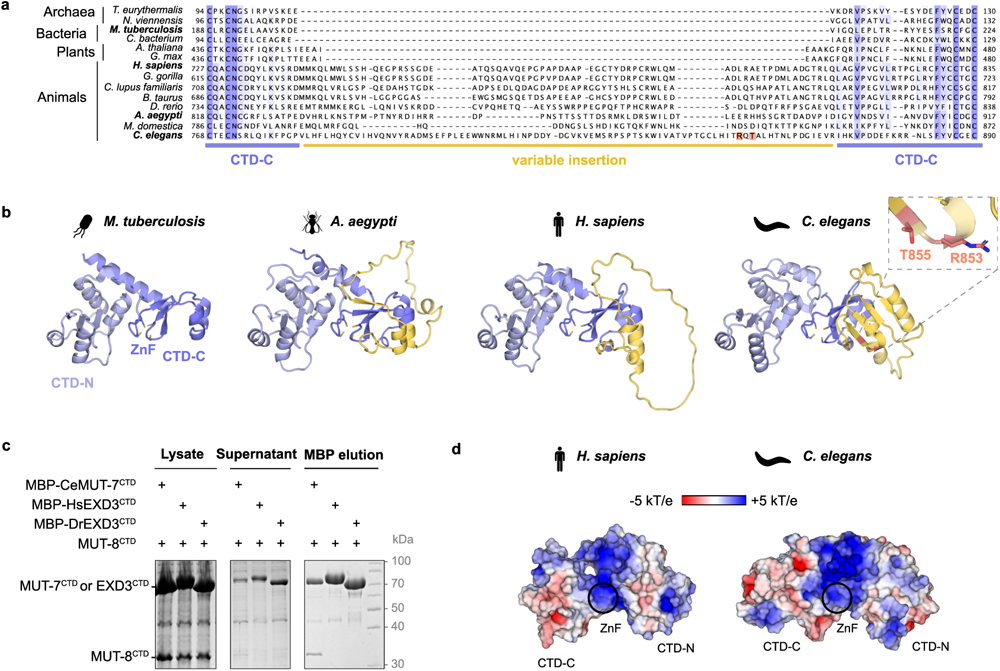
MUT7-C is an evolutionarily ancient domain. **a**, Multiple sequence alignment of bacterial, archaeal and eukaryotic MUT7-C domains, limited to the sequence between the first and second pair of cysteines of the zinc finger. Residues are marked with shades of blue based on their conservation score. The variable insertions are highlighted with a yellow line, MUT-7^CTD-C^ with a blue line. MUT-7^CTD^ residues preventing MUT-8 binding (R853, T855) are in orange. **b**, AlphaFold prediction of *M. tuberculosis* (O53776), *A. aegypti* (A0A7N4YH63), *H. sapiens* (Q8N9H8) MUT7-C domains and crystal structure of *C. elegans* MUT-7 CTD (PDB: 8Q66). The MUT-7^CTD-N^ is shown in light blue, the MUT-7^CTD-C^ in blue and the insertions in yellow. A zoom-in shows *C. elegans* MUT-7 mutated residues (R853, T855) in orange. **c**, Co-expression pulldown assays testing the species-specificity of the MUT-7^CTD^/MUT-8^CTD^ interaction. MBP-tagged CeMUT-7^CTD^, HsEXD3^CTD^ and DrEXD3^CTD^ were co-expressed in *E. coli* with CeMUT-8^CTD^. Hs= *Homo sapiens*; Dr= *Danio rerio*; Ce= *Caenorhabditis elegans*. After lysis, the supernatant was incubated with amylose resin. Total lysate, supernatant and elutions were analysed by SDS-PAGE followed by Coomassie staining. **d**, Electrostatic surface potential of HsEXD3^CTD^ (AF-Q8N9H8) and CeMUT-7^CTD^ (PDB: 8Q66). Electrostatic surface potential was calculated by Adaptive Poisson-Boltzmann Solver from −5 kT/e (red) to +5 kT/e (blue).

While the MUT-7^CTD-N^ domain is evolutionarily highly conserved at both the sequence and structural level (Fig. 6b and **Supplementary** Fig. 7a), MUT-7^CTD-C^ exhibits significant variability among species. Animals contain various insertions between the first and second pair of cysteines of the zinc finger (Fig. 6a and 6b). In most animals, the insertion is mainly disordered (e.g. in insects or vertebrates), but in *C. elegans* (and other Caenorhabditis species), it forms a four-stranded β-sheet (aa 784-868) (Fig. 6b). The surface of the β-sheet participates in MUT-8 binding and mutations in this region (R853E, T855E) abolish complex formation (Fig 4b and **6b**). Given the sequence and structural differences of nematode and vertebrate insertions, we hypothesised that the CTDs from *H. sapiens* (Hs) or *D. rerio* (Dr) EXD3 would not bind MUT-8. To test this, we co-expressed the MUT-8^CTD^ with MBP-tagged HsEXD3^CTD^ and DrEXD3^CTD^ and analysed the interaction by pulldown experiments. MUT-8^CTD^ was not soluble when co-expressed with HsEXD3^CTD^ or DrEXD3^CTD^, suggesting that the MUT-8^CTD^ does not interact with the vertebrate ortholog EXD3 and supporting the species-specificity of the MUT-7/MUT-8 interaction (Fig. 6c).

Our findings reveal that the evolutionarily conserved MUT7-C domain has acquired insertions that differ significantly across species. While the role of the insertion in other organisms remains unknown, the well-structured insertion present in Caenorhabditis mediates binding to MUT-8, thereby allowing recruitment of MUT-7 to Mutator foci.

### The MUT7-C domain has a conserved role in RNA-binding

What is the conserved, ancestral function of the MUT7-C domain? Mapping the electrostatic surface potential on the nematode and human MUT7-C domains, revealed a highly positively charged surface (Fig. 6d). Moreover, the MUT7-C domain contains a zinc finger, a motif often involved in nucleic acid binding. This prompted us to investigate the contribution of the MUT7-C domain to RNA binding for both worm MUT-7 and human EXD3 by fluorescence polarisation (FP) assays, using a 5′-FAM-labeled 16-mer ssRNA. For that, we purified the following constructs: MUT-7^FL^/MUT-8^CTD^, MUT-7^NTD-EXO^, MUT-7^CTD^/MUT-8^CTD^, catalytically inactive EXD3^FL(D399A)^ (1-876), EXD3^NTD-EXO(D399A)^ (1-582), and EXD3^CTD^ (624-876). We also verified the presence of Zn^2+^ in EXD3^CTD^ by SEC combined with inductively coupled plasma mass spectrometry (**Supplementary Fig.8a** and **8b**). MUT-7^FL^/MUT-8^CTD^ complex bound ssRNA with an affinity in the nanomolar range (*K*_d_ = 0.17 µM). MUT-7^NTD-EXO^, which lacks the CTDs of both MUT-7 and MUT-8, had a roughly 12-fold weaker affinity (*K*_d_ = 2.06 µM), while the isolated MUT-7^CTD^/MUT-8^CTD^ complex only bound RNA weakly (Fig. 7a and **Supplementary** Fig. 8c). To avoid precipitation of human EXD-3, experiments were performed at 250 mM NaCl, while experiments with *C. elegans* MUT-7 were done in a buffer containing 50 mM KCl. The results for EXD3 were similar to those obtained for MUT-7. The affinity of EXD3^FL(D399A)^ for the ssRNA was in the micromolar range (*K*_d_ = 1.86 µM). EXD3^NTD-EXO(D399A)^ had a roughly 6-fold weaker affinity (*K*_d_ = 11.15 µM), while the EXD3^CTD^ alone only bound RNA weakly (Fig. 7b and **Supplementary** Fig. 8d). To compare MUT-7 and EXD3 affinities for RNA, we tested RNA binding of a catalytically inactive MUT-7^FL(D435A)^/MUT-8^CTD^ complex at 250 mM NaCl, which almost completely abolished the binding of MUT-7 to RNA (Fig. 7b). This suggests that human EXD3 binds RNA more strongly than *C. elegans* MUT-7, which could be explained by the fact that the EXD3 NTD is more positively charged than its worm counterpart (**Supplementary** Fig. 8e).

**Figure 7:**
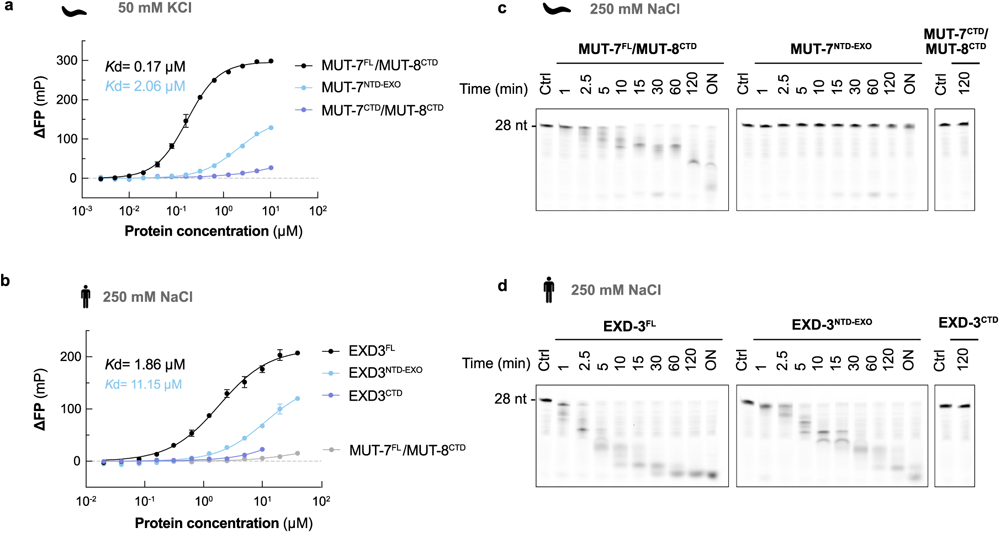
The MUT7-C contributes to the RNA binding properties of MUT-7 and EXD3. **a**-**b,** RNA-binding affinity of different MUT-7 and EXD3 constructs determined by Fluorescence Polarisation (FP) assays. A 5′-FAM-labelled 16-mer ssRNA (50 nM) was incubated with increasing concentrations of the indicated proteins. Fluorescence polarisation values were normalised by subtracting the FP values of the wells containing the labelled RNA alone. The mean of three experiments is shown, and error bars correspond to ± SD. The FP data are fitted to the Hill equation to obtain the dissociation constant, *K*d. RNA binding of MUT-7^FL^/MUT-8^CTD^, MUT-7^NTD-EXO^, and MUT-7^CTD^/MUT-8^CTD^ was tested in a buffer with 50 mM KCl and 5 mM EDTA (a). For EXD3^FL(D399A)^, EXD3^NTD-EXO(D399A)^, EXD3^CTD^ and MUT-7^FL(D435A)^/MUT-8^CTD^, a buffer containing 250 mM NaCl was used (b). **c**-**d**, Time-course assessing the nuclease activity of indicated MUT-7 (c) and EXD3 (d) constructs. Proteins (0.5 μM) were incubated with a 5′-FAM 28-mer ssRNA (1 μM) in a buffer containing 250 mM NaCl at room temperature for the indicated times. MUT-7^FL^/MUT-8^CTD^, MUT-7^NTD-EXO^, MUT-7^CTD^/MUT-8^CTD^, EXD3^FL^, EXD3^NTD-EXO,^ and EXD3^CTD^ were tested. Reaction products were separated by TBE-Urea PAGE.

Overall, our results indicate that for both worm MUT-7 and human EXD3, the MUT7-C domain synergistically contributes with the NTD and EXO domains to RNA binding. To investigate the functional contribution of the CTD to MUT-7/EXD3 nuclease activity, we compared the RNA degradation kinetics of MUT-7 and EXD3 in the presence and absence of the CTD. To this end, we performed a time course with a 5′-FAM-labeled 28-mer ssRNA using the following constructs: MUT-7^FL^/MUT-8^CTD^, MUT-7^NTD-EXO^, and the newly purified active EXD3^FL^ and EXD3^NTD-EXO^. The MUT-7^CTD^/MUT-8^CTD^ complex and EXD3^CTD^ were used as negative controls. For both MUT-7 and EXD3, the full-length constructs degraded RNA more efficiently than the ones lacking the CTD (Fig. 7c and 7d). EXD3 constructs degraded RNA with faster kinetics than their MUT-7 counterparts (Fig. 7c and 7d), consistent with the higher RNA binding affinity of EXD3 at 250 mM NaCl (Fig. 7b). Our findings suggest that the MUT7-C domain contributes to substrate recognition thereby improving the RNA degradation rate. This can likely be extended to other MUT-7 homologs.

## Discussion

In this study, we have structurally and functionally characterised the MUT7-C, an ancient domain belonging to the conserved 3’-5’ exoribonuclease MUT-7 in animals, while in prokaryotes it exists mostly as an individual protein.

Prokaryotes have a reduced, minimal version of the MUT-7C domain consisting of a TOPRIM-like fold and a zinc finger at the C-terminus. We did not detect significant structural similarity with PIN-like domains, as previously suggested, indicating that MUT7-C and PIN domains are only distantly related. Animals contain variable insertions within the zinc finger of the MUT7-C domain. Thus, the MUT7-C appears structurally plastic, suggesting a possible functional adaptation of this domain during evolution.

In Caenorhabditis, the MUT7-C insertion differs significantly from other animals at the sequence and structural level. Here, it forms a structured β-sheet, constituting the main binding site for the nematode-specific factor MUT-8. MUT-8 is a modular protein that binds MUT-7 and the Mutator scaffold MUT-16, acting as a bridge and recruiting MUT-7 to Mutator foci. The interaction between MUT-8 and MUT-7 is mediated by the two folded CTDs forming a tight complex. In contrast, the interaction between MUT-8 and MUT-16 is mediated by disordered regions of both proteins: the MUT-8 NTD and an internal MUT-16 segment. Mutations in the MUT7-C β-sheet prevent MUT-8 binding and result in RNAi resistance. Similarly, worm strains carrying nonsense mutations in MUT-7 CTD upstream of the insertion (*pk719*: W811/* and *pk204*: W812/*, see Figure 1a) show transposon de-silencing and RNAi resistance ^19^. These mutations result in truncations of the MUT-7 CTD, leading to loss of MUT-8 binding. This strongly suggests that correct localisation of MUT-7 at Mutator foci depends on MUT-8 binding and is necessary for sRNA amplification. Consistently, the MUT-7 paralog ZK1098.3 (P34603), which contains the NTD and EXO domains but only a partial CTD lacking the MUT-8 binding site, does not compensate for MUT-7 mutations.

But what is the role of MUT-7 in the sRNA amplification process at Mutator foci? The 3’-5’ exoribonuclease activity of MUT-7 is necessary for sRNA amplification, as *mut-7* mutants show a drop in 22G RNAs ^19,40^, and mutations close to (G497E) or in the catalytic site (E437K) lead to RNAi resistance and TE mobilisation ^19,24^.

Based on the current sRNA amplification model, the most likely hypothesis is that MUT-7 acts either on the pUG tail added by the nucleotidyltransferase MUT-2 or the RNA products synthesised by the RdRP RRF-1. We have shown that MUT-7 can bind unfolded single-stranded RNAs and the pUG fold with similar affinities (**Supplementary** Fig. 8f). However, MUT-7 only efficiently degrades unstructured RNAs. The pUG tail seems to serve two functions at a time: stabilising the newly formed 3’ end of the 5’ fragment and marking it as a template for the RdRP to initiate sRNA amplification ^17^. One could envisage that MUT-7 either trims the 3’ of RdRP products to their final length of 22 nt, similar to the function of Nibbler in Drosophila. Alternatively, MUT-7 could control the length of the pUG tail or even degrade abortive RdRP transcripts that are too short to function efficiently in RNAi. However, further studies are necessary to unravel the molecular function of MUT-7.

In the attempt to dissect the ancient function of the MUT7-C domain unrelated to the insertion, we showed that the CTD contributes to RNA binding for both *C. elegans* MUT-7 and the human homolog EXD3. Deletion of the CTD decreases the efficiency of RNA-binding and RNA degradation, suggesting that it is functionally important. This might indicate that RNA binding is an evolutionary feature of the MUT-7 CTD. While the ancient function of the MUT-7C domain in archaea and bacteria remains unknown, it is conceivable to speculate that it functions as an RNA binding module. This is supported by the observation that heterologous expression of *M. tuberculosis* MUT-7C homolog Rv0579 in *E. coli* leads to co-purification of nucleic acids ^41^. Dissecting the function of Rv0579 would be clinically relevant, as a mutation in Rv0579 results in resistance to the prodrug TP053 by an unidentified mechanism ^41^.

## Material and Methods

### Cloning and protein production

The genes coding for CeMUT-7 (P34607), CeMUT-8 (Q19672), CeMUT-16 (O62011), HsEXD3 (Q8N9H8) and DrEXD3 (A0A8M9Q6C2) as well as the respective truncations were cloned into modified pET vectors using ligation independent cloning. DrEXD3 and codon-optimized HsEXD3 cDNAs were synthesised by Twist Bioscience. Mutations were generated by the Quick-Change mutagenesis approach. Proteins were produced as fusion proteins with varying fusion tags in the *E. coli* BL21(DE3) derivatives strain in Terrific Broth (TB) medium. To reconstitute FL and truncated MUT-7/MUT-8 and MUT-8/MUT-16 complexes, MUT-7 was co-expressed with MUT-8 and MUT-8 with MUT-16, respectively. Protein production was induced at 18°C by adding 0.2 mM IPTG for 12-16 hours. An overview of the constructs used in this study and the purification steps used is provided in **Table S2**. Proteins were stored at −70°C, usually in 20 mM HEPES/NaOH (pH 7.5), 150 mM NaCl, 2 mM DTT, with or without 10% (v/v) glycerol. 0.5 mM TCEP was used as a reducing agent instead of the DTT for constructs containing the MUT7-C domain with the zinc finger. EXD3 constructs were stored in 20 mM HEPES/NaOH (pH 7.5), 500 mM NaCl, 0.5 mM TCEP.

### Co-expression pulldown assays

For interaction studies by the co-expression co-purification strategy, two plasmids containing the genes of interest and different antibiotic resistance markers were co-transformed into BL21(DE3) derivative strains to allow co-expression (**Table S3**). Cells were grown in 50 mL TB medium shaking at 37°C, and protein production was induced at 18 °C by adding 0.2 mM IPTG for 12-16 hours. Cell pellets were resuspended in 4 mL of lysis buffer (50 mM NaH_2_PO_4_, 20 mM Tris/HCl, 250 mM NaCl, 10 mM Imidazole, 10% (v/v) glycerol, 0.05% (v/v) IGEPAL, 5 mM 2-mercaptoethanol pH 8.0). Cells were lysed by sonication, and insoluble material was removed by centrifugation at 21,000xg for 10 minutes at 4°C. 500 µL supernatant was applied to 35 µL amylose resin (New England Biolabs) or glutathione resin (Cytiva) and incubated for 1-2 hours at 4°C. Subsequently, the resin was washed three times with 500 µL lysis buffer. The proteins were eluted in 50 µL of lysis buffer supplemented with 10 mM maltose (amylose resin) or 20 mM of reduced glutathione (glutathione resin), respectively. Input and eluate fractions were analysed by SDS–PAGE and Coomassie staining.

### Pulldown assays with purified proteins

To analyse protein interaction with purified proteins, the proteins (5 µM) (**Table S2**) were pre-incubated in 50 µL binding buffer containing 20 mM HEPES/NaOH (pH 7.5), 150 mM NaCl, 10% (v/v) glycerol, 0.05% (v/v) IGEPAL, 2 mM DTT for 30 minutes at 4°C. Samples were added to 20 µL glutathione resin (Cube Biotech) and gently shaken for 1-2 hours at 4°C. Beads were then washed three times with 200 μL binding buffer, and the retained material was eluted with 50 µL incubation buffer supplemented with 20 mM of reduced glutathione. Input material and one-third of the eluate were analysed by SDS-PAGE and Coomassie staining.

### Crystallisation

Crystallisation trials with the MUT-7^CTD(633–89^^9)^/MUT-8^CTD(322–56^^7)^ complex in SEC buffer (20 mM HEPES/NaOH pH 8.5, 150 mM NaCl, 0.5 mM TCEP) were performed using a vapour diffusion set-up by mixing the sample and crystallisation solution in a 1:1 and 2:1 ratio. Initial needle-shaped crystals were obtained at a protein concentration of 8 mg/ml in condition E3 (0.2 M NaI, 20% (w/v) PEG 3350) from the PACT screen (Molecular Dimensions). We obtained single, plate-shaped crystals through two rounds of cross-matrix micro-seeding using the Morpheus screen (Molecular Dimensions). The best crystals grew in condition A10 (10% (w/v) PEG 8000, 20% (v/v) ethylene glycol, 0.3 M MgCl_2_, 0.3 M CaCl_2_, 0.1 M bicine/Trizma base pH 8.5). Crystals from this condition did not require further cryoprotection and were, therefore, directly frozen in liquid nitrogen before data collection at 100 K.

### Data processing, phase determination, refinement, and model building

Diffraction data were processed automatically by the Grenoble Automatic Data Processing (GrenADES) pipeline from ESRF (https://www.esrf.fr/UsersAndScience/Experiments/MX/How_to_use_our_beamlines/ Run_Your_Experiment/automatic-data-processing). Here, data were integrated with XDS ^42^ and further processed using pointless and aimless ^43,44^. The phases were determined by molecular replacement using the AlphaFold model of *C. elegans* MUT-7 C-terminal domain AF-P34607-F1 (residues 639-898) (https://alphafold.ebi.ac.uk/entry/P34607). Molecular replacement was performed with Phaser ^45^ within Phenix ^46^. Preceding molecular replacement, the model was prepared with Phenix (process predicted model) to translate the pLDDT values to B factors and to remove flexible regions. Following molecular replacement, the model was automatically built with ModelCraft ^47,48^ within CCP4i2 ^49^, manually completed with COOT ^50^ and refined with phenix.refine ^51^ and refmac5 ^52^. Model quality was assessed using molprobity ^53^ and PDB-REDO ^54^. Data collection and refinement statistics are shown in **Table S1**. Molecular graphics of the structures were created using PyMOL 2.5.2.

### Ribonuclease activity assays

RNAs carrying a 5′-6-fluorescein amidite (5′-FAM) label were purchased from Ella Biotech (Fuerstenfeldbruck, Germany): a 28-mer ssRNA (AUUGCAUCUAAAGUUGAUUGAAGAGUUC), a p(GU)_14_ ssRNA (GUGUGUGUGUGUGUGUGUGUGUGUGUGU) and a p(GU)_8_ ssRNA (GUGUGUGUGUGUGUGU). Lyophilised RNAs were resuspended in H_2_O at a final concentration of 100 µM and stored at −70 °C. To fold the two p(GU) RNAs, RNAs were diluted in folding buffer (50 mM Tris/HCl pH 7.5, 100 mM KCl) at a final concentration of 20 µM; samples were heated at 90°C for 3 minutes and then slowly cooled down decreasing the temperature by 1 °C/minute to 4°C. Different concentrations of recombinant proteins were incubated with 1 µM RNA for 30 minutes at room temperature in a total volume of 10 µL in a buffer containing 20 mM HEPES/NaOH, 50 mM KCl and 2 mM MnCl_2_ unless stated otherwise. For the cation-dependent activity assay, the buffer contained 5 mM of different divalent cations as indicated. For the pUG RNA degradation assay, the buffer contained 50 mM KCl or LiCl as indicated. For the time course experiments comparing *C. elegans* MUT-7 and *H. sapiens* EXD3 activities, a buffer with 20 mM HEPES/NaOH, 250 mM NaCl and 2 mM MnCl_2_ was used. To stop the reactions, 10 μL Gel Loading Buffer II (Invitrogen) was added, and samples were incubated at 98 °C for 5 minutes. Reaction products were resolved on homemade 15% TBE-Urea Gel. Gels were scanned with a Typhoon FLA-9500 imager (GE Healthcare).

### Fluorescence polarisation (FP) experiments

RNAs carrying a 5′-6-fluorescein amidite (5′-FAM) label were purchased from Ella Biotech (Fuerstenfeldbruck, Germany): a 16-mer ssRNA (GUUGAUUGAAGAGUUC) and a p(GU)_14_ ssRNA (GUGUGUGUGUGUGUGUGUGUGUGUGUGU). Lyophilised RNAs were resuspended in H_2_O at a final concentration of 100 µM and stored at −70°C. The p(GU)_14_ RNA was folded as described above. Increasing concentrations of proteins (from 0.002 to 10 µM) were incubated with 50 nM of RNA in a volume of 20 µL at room temperature for 30 minutes. For the *C. elegans* proteins, a buffer containing 20 mM HEPES/NaOH (pH 7.5), 50 mM KCl, and 5 mM EDTA was used unless stated otherwise. For the *H. sapiens* proteins, the buffer contained 20 mM HEPES/NaOH (pH 7.5) and 250 mM NaCl. The fluorescence polarization data were recorded on a Tecan SPARK plate reader with the following settings: excitation 485 nm, bandwidth 20 nm; Emission 535 nm, bandwidth 25 nm. Milli-polarization (mP) values were normalised by subtracting the mP values of the wells containing the fluorophore-labelled RNA only. Data were analysed using the nonlinear regression Hill equation within GraphPad Prism version 10. The mean of three replicates ± SD is shown.

### Circular dichroism (CD)

Far-UV CD spectra were measured using a Chirascan plus spectrometer (Applied Photophysics) and a 0.5-mm path-length cuvette at 20°C. Samples were prepared at 7.8 μM in 5 mM NaH_2_PO_4_, 80 mM NaF at pH 8.0. Eight spectra between 190-260 nm (0.5 nm steps) were collected and averaged. After subtraction of the buffer spectrum and conversion of ellipticity (θ) to mean residue ellipticity (MRE), values were plotted using GraphPad Prism version 10.0.2.

### In vitro condensate formation assays

Proteins were diluted to a final concentration of 10 µM in 20 mM Tris/HCl (pH7.5), 150 mM NaCl, 2 mM DTT and 5% (w/v) PEG6000. MUT-7/MUT-8 was mixed with MUT-16^584–7^^24^. 50 µL were immediately loaded onto a 96-well Greiner sensoplate pre-coated with 1% Pluronic F-127. The plate was incubated on the microscope for 100 minutes to allow droplet formation. Wells were imaged with a Zeiss Axio Observer Z1 in bright field mode, EC Plan-Neofluar 100x/1.3 Oil M27, Orca Flash 4.0 LT+ Camera, VIS-LED at 50 % intensity and 20 ms exposure time. Images were analysed with Fiji/ImageJ ^55^. Scale bars correspond to 50 µm.

### Multiple sequence alignment (MSA)

The following proteins were selected for the sequence alignment of the MUT7-C domains: *Thermococcus eurythermalis* (A0A097QWN3/1-153) Mut7-C RNAse domain-containing protein; *Nitrososphaera viennensis* (A0A060HM51/1-171) Mut7-C RNAse domain-containing protein; *Mycobacterium tuberculosis* (O53776/1-252) Twitching motility protein PilT; *Chlamydiota bacterium* (A0A7X5Q8E0/1-151) Mut7-C RNAse domain-containing protein; *Arabidopsis thaliana* (A0A1P8BGD5/1-516) 3’-5’ exonuclease domain-containing protein; *Glycine max* (A0A0R0GNL3/1-505) 3’-5’ exonuclease domain-containing protein; *Homo sapiens* (Q8N9H8/1-876) Exonuclease mut-7 homolog; *Gorilla gorilla gorilla* (G3S583/1-764) Exonuclease 3’-5’ domain containing 3; *Canis lupus familiaris* (A0A8P0SNB1/1-905) Exonuclease 3’-5’ domain containing 3; *Danio rerio* (A0A8M9Q6C2/1-861) Exonuclease 3’-5’ domain-containing 3; *Aedes aegypti* (A0A7N4YH63/1-944) Uncharacterized protein; *Musca domestica* (A0A1I8NEK9/1-911) Exonuclease mut-7 homolog; *Caenorhabditis elegans* (P34607/1-910) Exonuclease mut-7. The MSA was performed with ClustalO ^56^ using standard parameters and was visualized with Jalview ^57^.

### Strain maintenance

Worm strains were cultured according to standard laboratory conditions at 20 °C on nematode growth medium (NGM) plates seeded with E. coli OP50 ^58^. The following strains were used in this study: wild type N2, *mut-7*(*pk204*) III (RFK502), *mut-7*(xf367[*mut-7*(R853E, T855E)]) III (RFK1714), *mut-16*(cmp3[*mut-16*::gfp::flag + loxP]) I (RFK1713), *mut-16*(cmp3[*mut-16*::gfp::flag + loxP]) I; *mut-7*(xf367[*mut-7*(R853E, T855E)]) III (RFK1716).

### Generation of mutant line

CRISPR gRNAs were designed using Integrated DNA Technologies CRISPR–Cas9 guide RNA design tool. Strains were obtained by injecting a recombinant Cas9 protein (homemade), a single-stranded primer as a template for homologous recombination (IDT) and a guide RNA molecule (IDT) as described previously in ^59^. The mutants were analysed by sequencing and outcrossed two times to wild-type N2 worms to remove all potential off-target. For generating the *mut-7* (xf367[*mut-7*(R853E, T855E)]) III (RFK1714) strain, the guide AACCTAATTTGGCTTTATTTAGG and repair template TTCAATTCCATCTGGCAGATTGGTGTGAAGTGCCTCTTGCTCTCTAAATAAAGCCAAATTAGGTTTTAATAATATT were used.

### RNAi

Adult animals were bleached to obtain embryos, which then hatched and synchronized in M9 buffer (22 mM KH_2_PO_4_, 42 mM Na_2_HPO_4_, 86 mM NaCl, 1 mM MgSO_4_) for 24 hours at 20°C. L1 animals were fed with bacteria carrying an empty vector (L4440) or a vector expressing dsRNA against *pos-1* that came from Ahringer library. Young adult worms were transferred to fresh RNAi plates for the egg-laying assay, remaining continuously exposed to RNAi bacteria. All embryos within the first 24 h of egg-laying were scored for hatching. The percentage of viable progeny (number of hatched larvae/total number of laid eggs) was calculated. The progeny of around 15-20 worms was analysed for each strain.

### Microscopy

For live imaging, 20–25 young adult worms were picked to a drop (80 μl) of M9 buffer (22 mM KH_2_PO_4_, 42 mM Na_2_HPO_4_, 86 mM NaCl, 1 mM MgSO_4_) on a slide and washed in M9 with 0.05M NaN_3_ to paralyse the worms. After removing M9, a slide prepared with 2% agarose (in water) was placed on top of the coverslip and worms were imaged directly. Images were acquired at a Leica TCS SP5 STED CW confocal microscope (objective HC PL APO CS2 40x 1.3 oil-immersion objective, Leica). Images were processed with Leica LAS software and ImageJ.

## Data availability

Coordinates and structure factors of the MUT-7^CTD^/MUT-8^CTD^ complex structure have been deposited in the Protein Data Bank (PDB) with accession code PDB ID 8Q66 (https://doi.org/10.2210/pdb8Q66/pdb).

## Acknowledgements

We thank all members of the Falk laboratory for fruitful discussions. Andrea Pauli and Victoria Deneke for discussions and support on EXD-3. Svenja Helmann for technical assistance on genome editing. Harald Hornegger for helping with the condensate formation assays. Andreas Brandstätter (BOKU, Vienna) for performing the SEC-ICP-MS measurements. Maria Novatchkova (IMP, Vienna) for phylogenetic analysis. All members of the VBC proteomics facility for their support.

## Author contributions

V.B. and S.F. conceived the study; V.B. designed, performed, and analysed experiments; L.P. performed and analysed genetic experiments in worms; R.L. contributed to protein purification; V.B. and S.F. solved the crystal structure; S.H. supervised SEC-ICP-MS measurements; S.F. supervised the study; R.F.K. assisted in data analysis and interpretation; V.B. made the figures; V.B. and S.F. wrote the manuscript with contributions from R.F.K..

## Supplementary information

### Supplementary Figures

**Supplementary Fig. 1:**
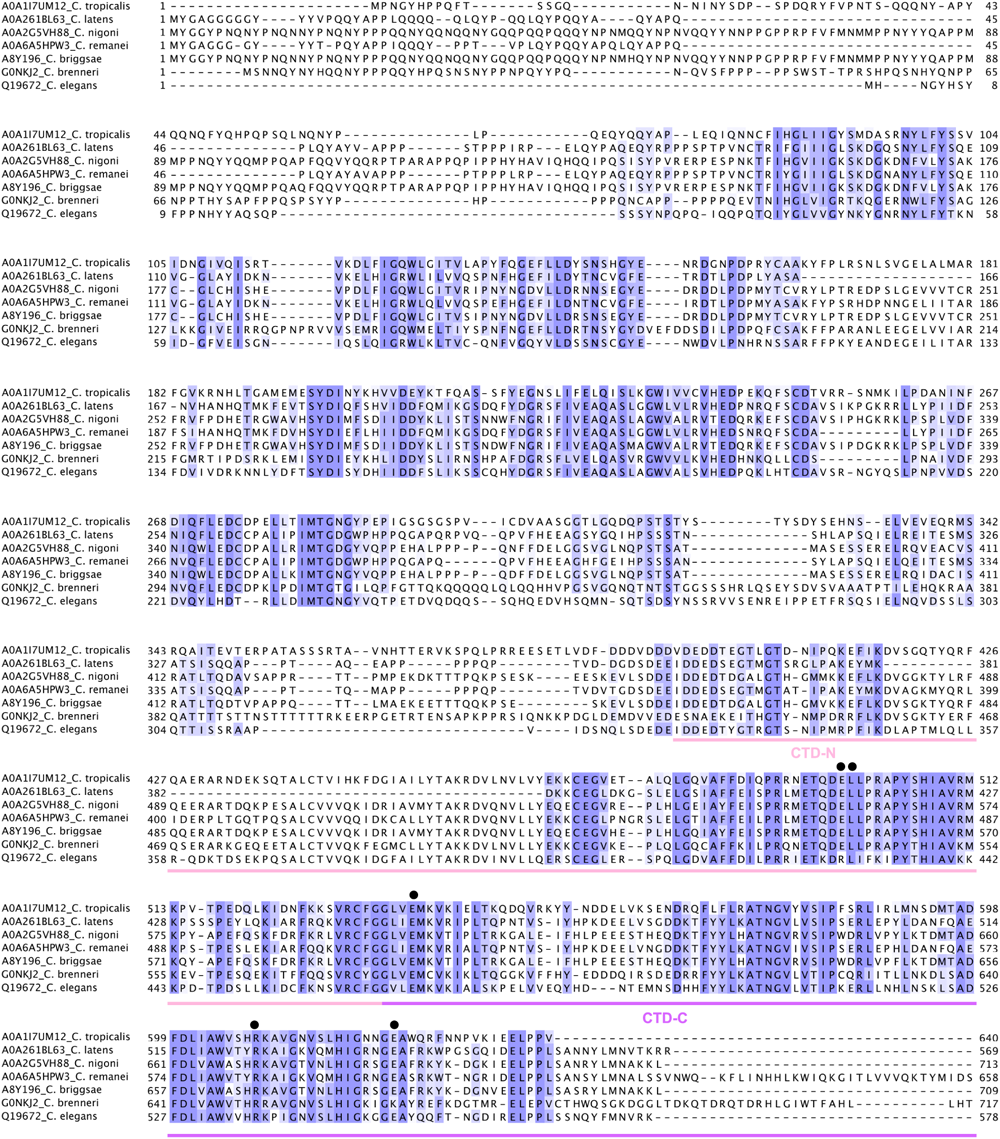
Multiple sequence alignment of different Caenorhabditis MUT-8 homologs. Residues are marked with shades of blue based on their conservation score. MUT-8 CTD subdomains are indicated with a light pink (CTD-N) or pink (CTD-C) line. MUT-8 residues involved in MUT-7 binding according to the crystal structure (PDB: 8Q66) are highlighted by black dots.

**Supplementary Fig. 2:**
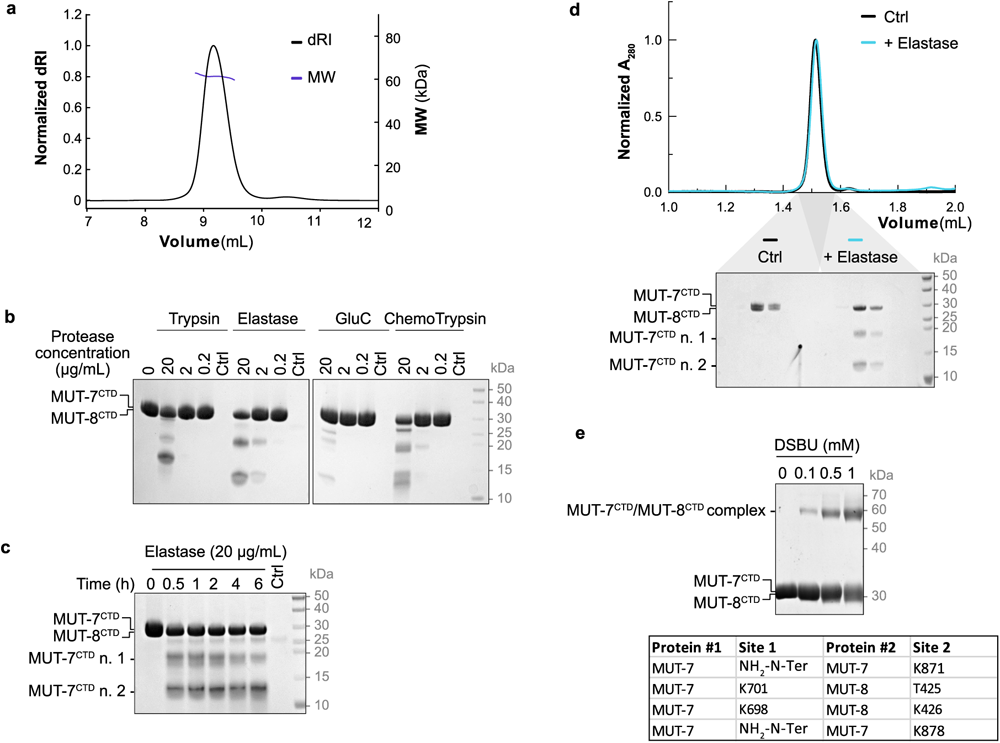
The MUT-7^CTD^/MUT-8^CTD^ complex is a stable heterodimer. **a**, SEC-Multi-Angle Light Scattering (SEC-MALS) analysis of the MUT-7^CTD^/MUT-8^CTD^ complex. The sample was loaded on a Superdex 75 increase (10/300) column. Normalised differential Refractive Index (dRI) and calculated Molecular Weight (MW) are shown. **b**, Limited proteolysis of the purified MUT-7^CTD^/MUT-8^CTD^ complex. MUT-7^CTD^/MUT-8^CTD^ was incubated with decreasing concentrations of indicated proteases for 30 minutes on ice. The control (Ctrl) sample corresponds to the protease alone at the highest concentration. Reaction products were analysed by SDS-PAGE followed by Coomassie staining. MUT-7^CTD^ has a theoretical MW of 32 kDa and MUT-8^CTD^ of 29 kDa, being indistinguishable on the gel where they appear as a single band. **c**, Time-course of the limited proteolysis assay of the MUT-7^CTD^/MUT-8^CTD^ complex. MUT-7^CTD^/MUT-8^CTD^ was incubated with elastase (20 µg/mL) on ice for the indicated amounts of time. The control (Ctrl) sample corresponds to the elastase alone. A control reaction without protease was also performed. Reaction products were analysed by SDS-PAGE followed by Coomassie staining. **d**, Size exclusion chromatography analysis of the native (black, Ctrl) and elastase-treated (teal, + Elastase) MUT-7^CTD^/MUT-8^CTD^ complex. The peak fractions were analysed by SDS-PAGE followed by Coomassie-staining. Upon elastase treatment, MUT-7^CTD^ (32 kDa) is cleaved into two fragments (MUT-7^CTD^ n.1 and MUT-7^CTD^ n.2), while MUT-8^CTD^ (29 kDa) remains intact. **e**, Crosslinking-MS analysis of the MUT-7^CTD^/MUT-8^CTD^ complex. MUT-7^CTD^/MUT-8^CTD^ was incubated with increasing concentrations of disuccinimidyldibutyric urea (DSBU). Samples were visualised by SDS-PAGE followed by Coomassie staining. The band at 60 kDa corresponding to the crosslinked MUT-7^CTD^/MUT-8^CTD^ heterodimer was cut and analysed by MS. The table illustrates identified crosslinked residues.

**Supplementary Fig. 3:**
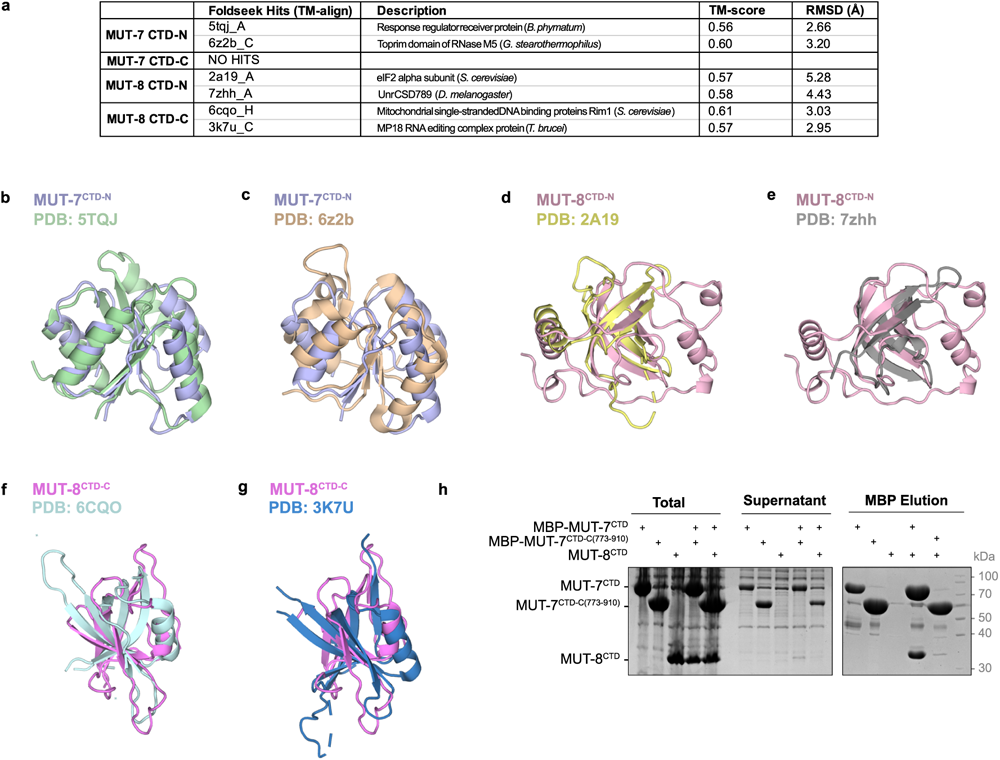
Structural analysis of MUT-7 CTD and MUT-8 CTD. **a**, Table with structures identified by Foldseek to be similar to MUT-7 CTD and MUT-8 CTD subdomains (CTD-N and CTD-C). TM-score and RMSD are reported. **b-g**, Structural alignments listed in (a). **b** and **c**, Structural alignment of MUT-7 CTD-N (light blue) with the response regulator receiver protein from *B. phymatum* (green) ^60^ (b) and the TOPRIM domain from *G. stearothermophilus* RNase M5 (wheat) ^6^^1^ (c). **d** and **e**, Structural alignment of MUT-8 CTD-N (light pink) with the *S. cerevisiae* eukaryotic translation initiation factor 2 alpha subunit (yellow) ^6^^2^ (d) and the *D. melanogaster* C-terminal cold shock domain of Upstream of N-Ras (grey) ^6^^3^ (e). **f** and **g**, Structural alignment of MUT-8 CTD-C(magenta) with the *S. cerevisiae* mitochondrial single-stranded DNA binding protein Rim1 (cyan) ^64^ (f) and the *T. brucei* MP18 RNA editing complex protein (blue) ^6^^5^ (g). **h**, Co-expression pulldown assay testing the interaction between two MBP-tagged MUT-7 constructs (MUT-7^CTD^ and MUT-7^CTD-C(773–910)^) and MUT-8^CTD^. MUT-7 constructs were co-expressed with MUT-8^CTD^ in *E. coli*. Expression of MUT-8^CTD^ alone was used as negative control. After lysis, the supernatant was incubated with amylose resin. Total lysate, supernatant and elution were analysed by SDS-PAGE followed by Coomassie staining.

**Supplementary Fig. 4:**
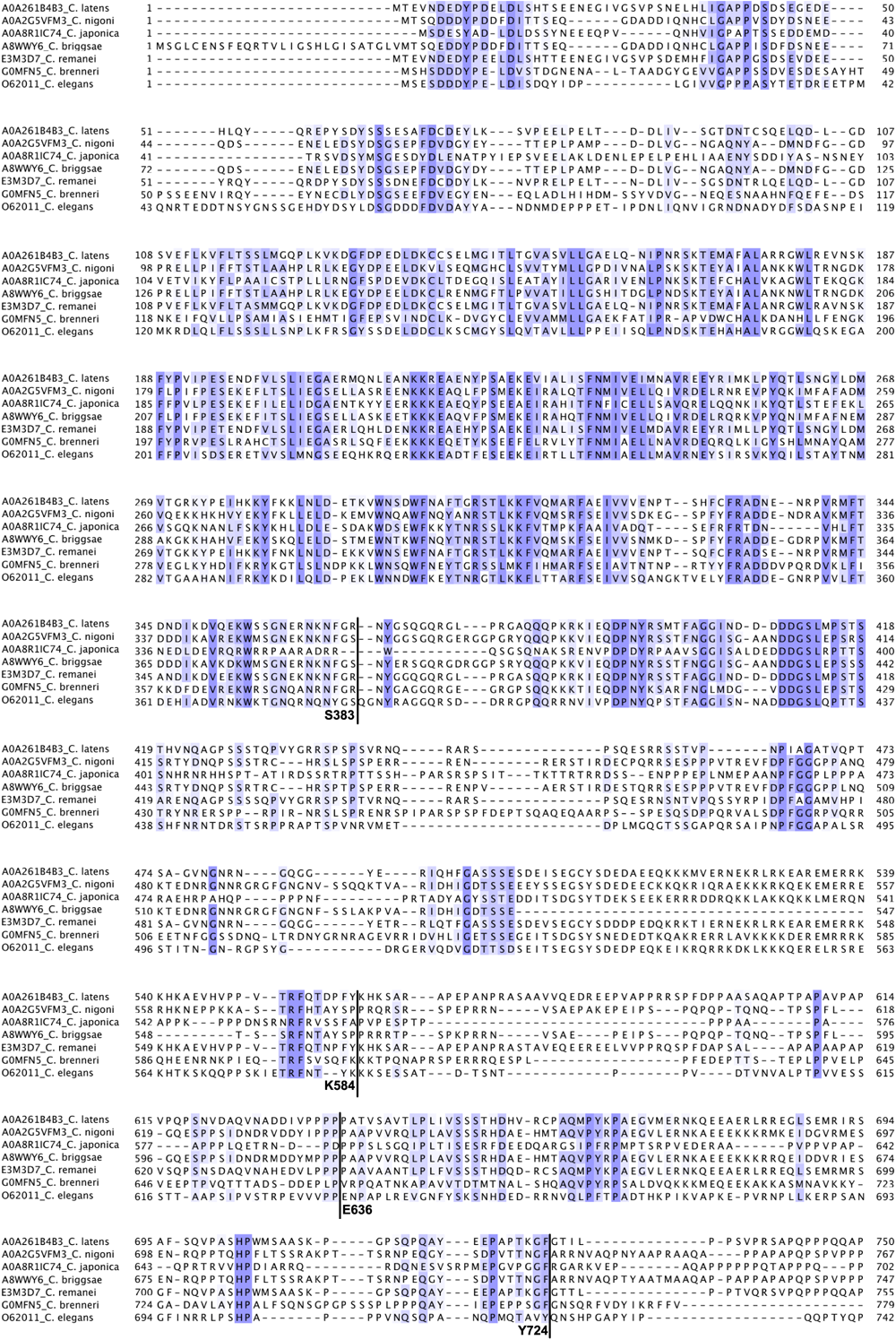
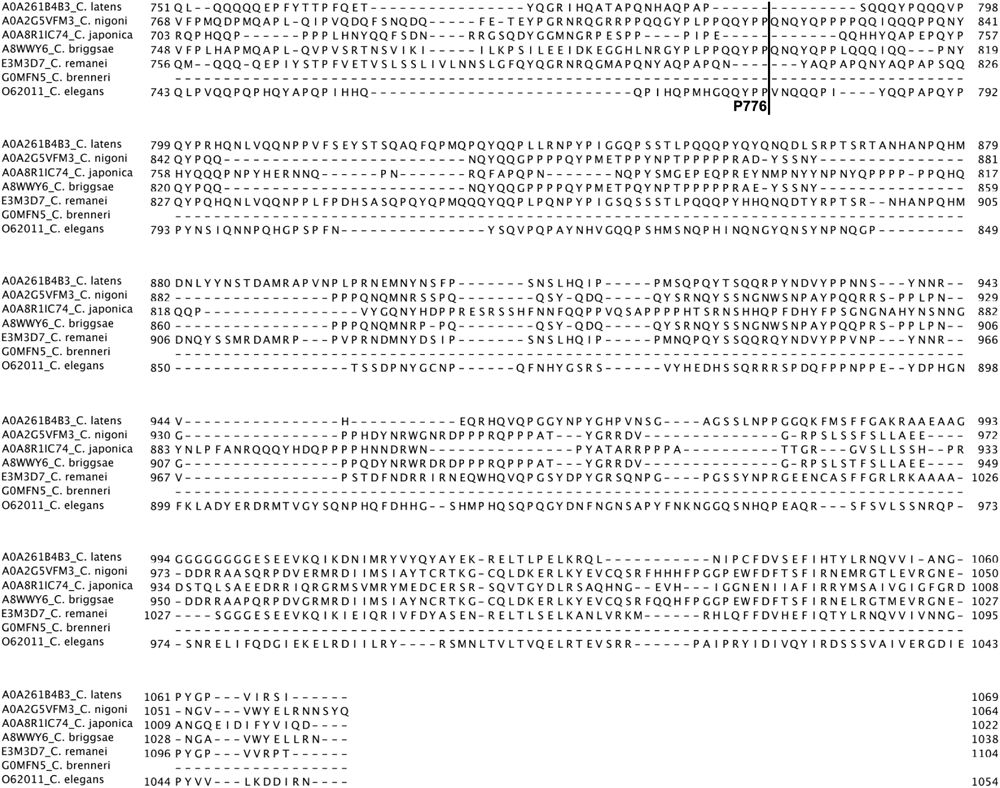
Multiple sequence alignment of different Caenorhabditis MUT-16 homologs. Residues are marked with shades of blue based on their conservation score. Boundaries of MUT-16 constructs mentioned in the text are highlighted.

**Supplementary Fig. 5:**
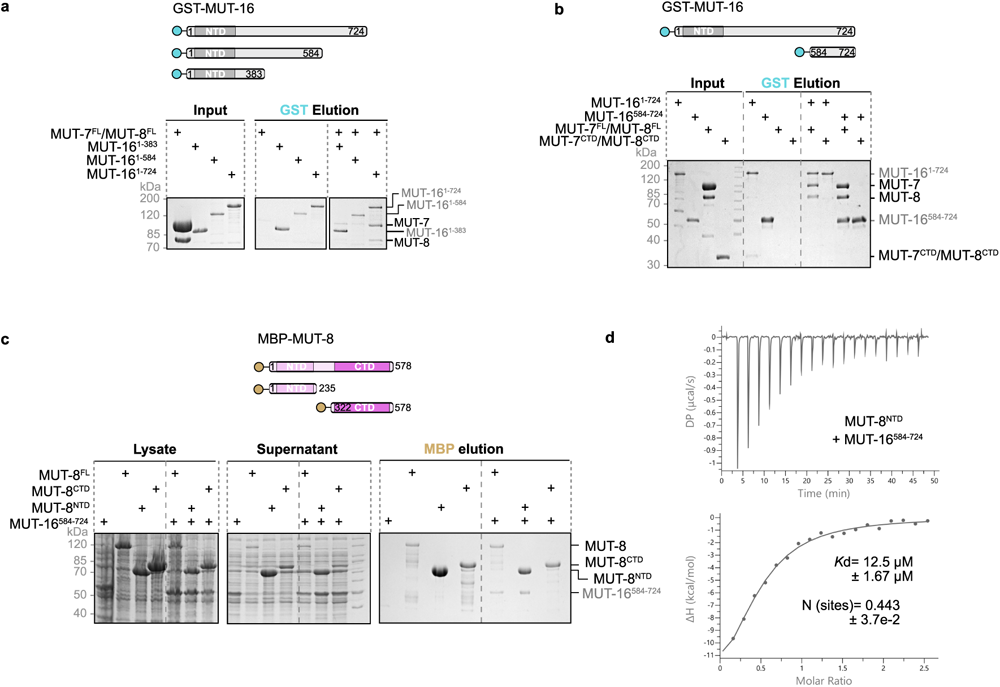
MUT-8 directly interacts with MUT-16. **a**, GST pulldown assessing the binding of three different GST-tagged MUT-16 constructs (MUT-16^1–724^, MUT-16^1–584^ and MUT-16^1–38^^3^) to the untagged MUT-7/MUT-8 complex. The cartoon illustrates the three MUT-16 constructs used as bait. Purified proteins were incubated with glutathione-coupled beads. Incubation of untagged MUT-7/MUT-8 with glutathione-coupled beads was used as negative control. Input and elution fractions were analysed by SDS-PAGE followed by Coomassie staining. **b**, GST pulldown assessing the binding of two GST-tagged MUT-16 constructs (MUT-16^1–724^ and MUT-16^584–724^) to the untagged MUT-7/MUT-8 and MUT-7^CTD^/MUT-8^CTD^ complexes. The cartoon illustrates the two MUT-16 constructs used as bait. Purified proteins were incubated with glutathione-coupled beads. Incubation of untagged MUT-7/MUT-8 and MUT-7^CTD^/MUT-8^CTD^ complexes with glutathione-coupled beads was used as negative control. Input and elution fractions were analysed by SDS-PAGE followed by Coomassie staining. **c**, Co-expression pulldown assay testing the interaction between MBP-tagged MUT-8 constructs (MUT-8, MUT-8^NTD^ and MUT-8^CTD^) and GST-tagged MUT-16^584–724^. The cartoon illustrates the three MUT-8 constructs used as bait. MUT-8 constructs were co-expressed with MUT-16^584–724^ in *E. coli*. Expression of GST-tagged MUT-16^584–724^ alone was used as negative control. After lysis, the supernatant was incubated with amylose resin. Total lysate, supernatant and elution are analysed by SDS-PAGE followed by Coomassie staining. **d**, Isothermal titration calorimetry (ITC) experiment analysing the interaction between MUT-8^NTD^ and MUT-16^584–724^. One representative of three ITC experiments is shown. The calculated dissociation constant (*K*d) and the number of binding sites (N) are the mean of three experiments, and the error bars correspond to ± SD. We note that the stoichiometry determined by ITC deviates from the expected 1 to 1 ratio.

**Supplementary Fig. 6:**
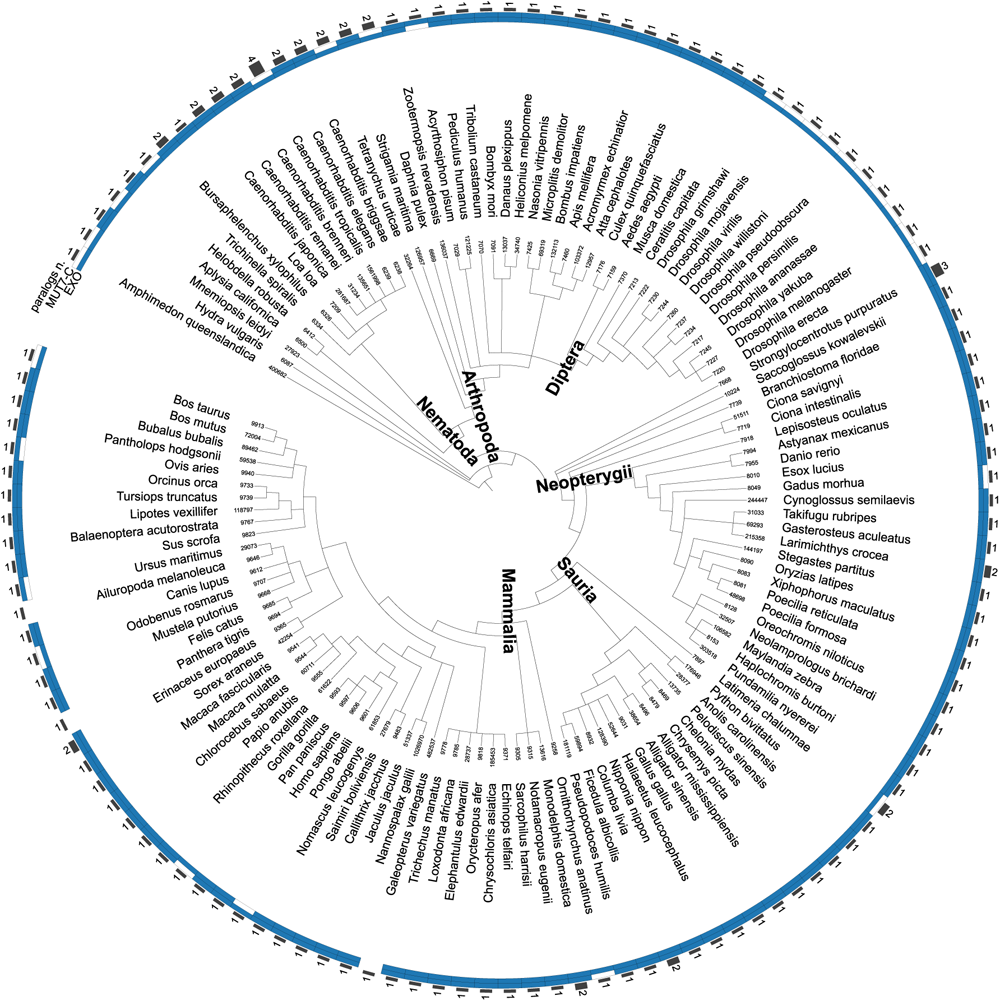
Phylogenetic species tree illustrating the conservation of MUT-7 and its CTD across metazoan. The tree is based on the EggNOG non-supervised orthologous group ENOG503BGHH (taxonomy level metazoan). The presence of MUT-7 exoribonuclease domain (PF01612) and its C-terminal domain (PF01927) in each species is marked in blue. The number of predicted homologs per species is also indicated.

**Supplementary Fig. 7:**
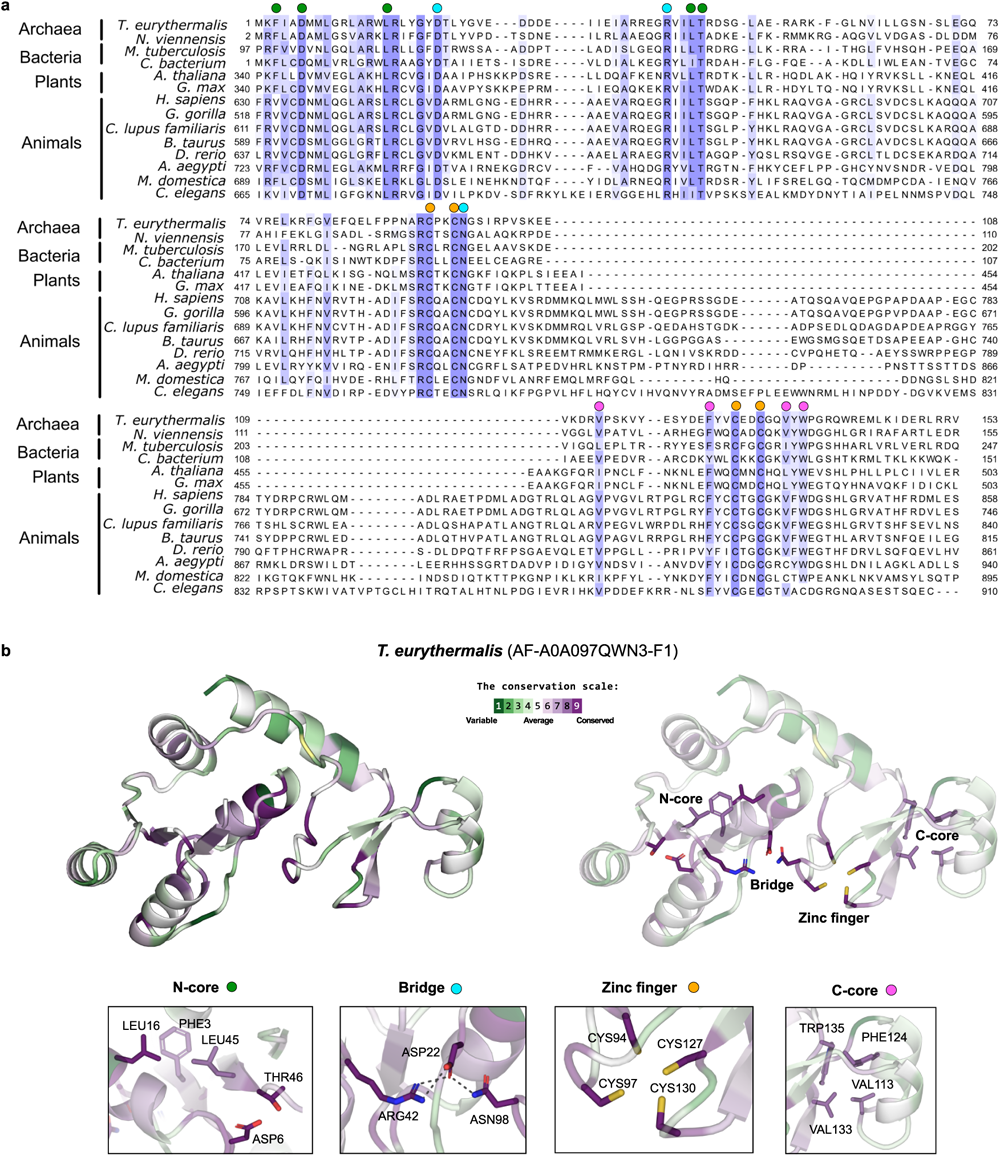
MUT7-C domain conservation. **a**, Multiple sequence alignment of bacterial, archaeal and eukaryotic MUT7-C domains. The conserved residues described in (b) are highlighted in green (for the N-core), cyan (for the Bridge), orange (for the Zinc finger) and magenta (for the C-core). **b**, Cartoon representation of the MUT7-C domain-containing protein A0A097QWN3 from the archaeon *Thermococcus eurythermalis* (AlphaFold prediction) coloured according to conservation. Conservation scores were calculated using ConSurf, with the sequences shown in (a). The gradient from green to purple indicates increasingly conserved residues. MUT-7^CTD-N^ contains a cluster of hydrophobic and charged residues that build the N-terminal core (N-core). The MUT-7^CTD-C^ is characterised by a zinc finger formed by the four invariant cysteine residues (Zinc finger) and a small core of hydrophobic and aromatic residues that flank the two C-terminal Cysteines of the zinc-ribbon (C-core). MUT-7^CTD-C^ is connected to MUT-7^CTD-N^ via a bridge element (Bridge) that is formed by the MUT-7^CTD-C^ Asn98 flanking the first two zinc-ribbon Cysteines and Arg42 and Asp22 in MUT-7^CTD-N^.

**Supplementary Fig. 8:**
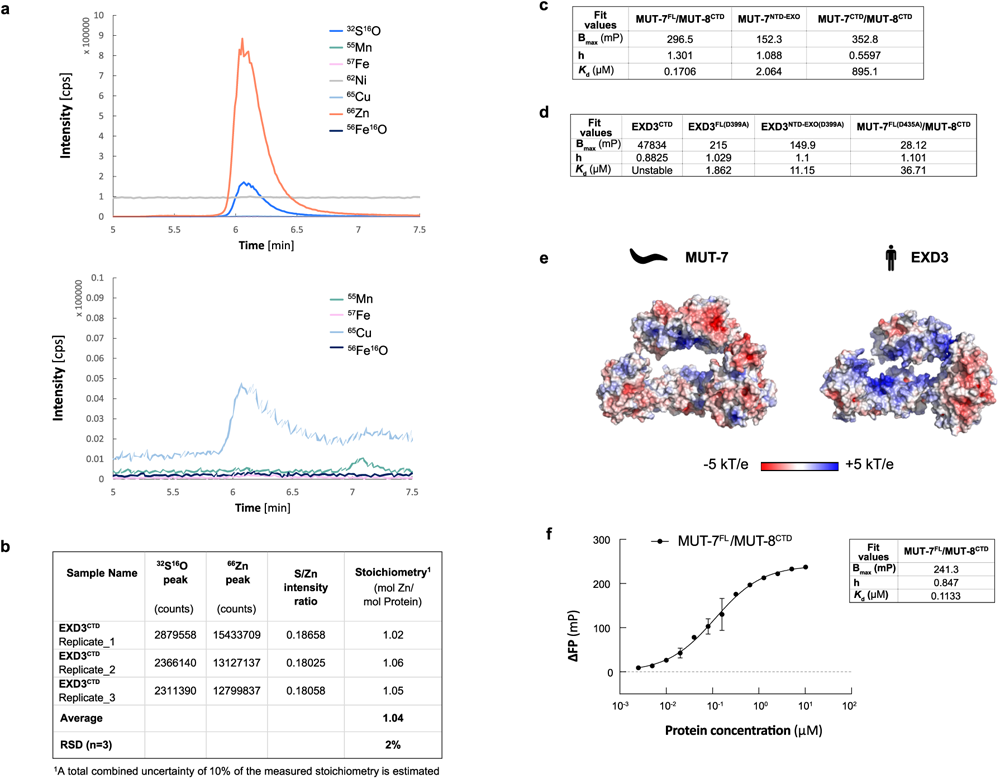
MUT-7 and EXD3 CTDs contribute to RNA binding. **a** and **b**, SEC-ICP-MS of human EXD3 CTD. **a**, Chromatograms showing the count per second (cps) obtained over time for different species. A zoom-in is shown in the lower panel. Data are representative of three technical replicates. **b**, Table summarizing the count per second (cps) values obtained for ^66^Zn and ^32^S^16^O in each of the three replicates. The intensity ratio between the ^66^Zn and the ^32^S^16^O signals was calculated. Comparison of this ratio with the value obtained for the calibrant (Bovine Cu/Zn-superoxide dismutase) allowed the determination of the number of Zinc ions bound to each EXD-3 molecule. **c**, Fit values for maximum specific binding value (Bmax), hill coefficient (h) and dissociation constant (*K*d) of the experiment testing MUT-7 affinity for a 16-mer RNA (see Fig. 7a). **d**, Fit values for maximum specific binding value (Bmax), hill coefficient (h) and dissociation constant (*K*d) of the experiment testing EXD3 affinity for a 16-mer RNA (see Fig. 7b) **e**, Electrostatic surface potential of MUT-7 and EXD3 AlphaFold predictions. Electrostatic surface potential was calculated by Adaptive Poisson-Boltzmann Solver (APBS) from −5 kT/e (red) to +5 kT/e (blue). **e**, pUG RNA-binding affinity of MUT-7^FL^/MUT-8^CTD^ determined by FP assays. A pre-folded 5′-FAM-labeled (GU)14 (50 nM) was incubated with increasing concentrations of protein in a buffer with 50 mM KCl and 5 mM EDTA. Fluorescence polarisation values were normalised by subtracting the FP values of the wells containing the labelled RNA alone. The mean of three experiments is shown with error bars corresponding to ± SD. The dissociation constant (*K*d), the hill coefficient (h) and the maximum specific binding value (Bmax) are derived by fitting the FP data to the Hill equation.

## Supplementary Tables

**Table S1:** Data collection and refinement statistics Rmsd: Root mean square deviation. https://doi.org/10.2210/pdb8Q66/pdb.

|  |  |
| --- | --- |
| Dataset | **MUT-7<sup>CTD</sup>/MUT-8<sup>CTD</sup>** |
| PDB | 8Q66 |
| Space group | C 1 2 1 |
| Cell dimensions |  |
| *a*, *b*, *c* (Å) | 145.09, 59.05, 88.16 |
| $\alpha$, $\beta$, $\gamma$ (°) | 90.0, 119.8, 90.0 |
| **Data Collection** |  |
| Wavelength (Å) | 0.87 |
| Resolution range | 67.91 – 2.03 (2.10 – 2.03) |
| No. of reflections | 146,924 (15006) |
| No. of unique reflection | 42,083 (4152) |
| *R*<sub>merge</sub> (%) | 8.9 (127.2) |
| *R*<sub>pim</sub> (%) | 5.6 (100) |
| *I* / σ*I* | 9.1 (0.8) |
| Completeness (%) | 99.7 (99.6) |
| Multiplicity | 3.5 (3.6) |
| CC<sub>1/2</sub> | 1 (0.41) |
| **Refinement** |  |
| Resolution range | 62.95 – 2.03 (2.10 – 2.03) |
| Reflections used in refinement | 41,976 (4145) |
| *R*<sub>work</sub> / *R*<sub>free</sub> (%) | 18.8 / 22.1 |
| Wilson B-factor (Å<sup>2</sup>) | 40.55 |
| Average B-factors (Å<sup>2</sup>) | 49.52 |
| **No. of atoms** |  |
| Proteins | 4168 |
| Ligands | 31 |
| Solvent | 250 |
| **Stereochemistry** |  |
| Rmsd bond lengths (Å) | 0.0103 |
| Rmsd bond angles (°) | 0.56 |
| Ramachandran favored (%) | 97.8 |
| Ramachandran outliers (%) | 0.2 |

**Table S2:** Recombinant proteins and purification strategies used in this study. “/” indicates that the two proteins were co-expressed and purified as a complex. (N) and (C) indicate the location of the tag at either the N-or C-terminus. His10 or His6 indicate the presence of 10 or 6 Histidine residues. MBP = Maltose Binding Protein, Trx = Thioredoxin, GST = Glutathione S-transferase, 3C = 3C cleavage site (LEVLFQ/GP), SEC = Size Exclusion Chromatography, SEC-MALS = Size Exclusion Chromatography (SEC) coupled to multi-angle light scattering (MALS), CD= Circular dichroism.

| Name | Residues | Tags | Purification Steps | Plasmid # | Use |
| --- | --- | --- | --- | --- | --- |
| MUT-7<sup>FL</sup> | 1-910 | (N) His10-MBP-3C | MBPTrap, 3C, Heparin, SEC | Ce255 | Nuclease assay |
| MUT-7<sup>NTD-EXO</sup> | 1-625 | (N) His10-MBP-3C | HISTrap, 3C, HISTrap, Heparin, SEC | Ce333 | Nuclease assay |
| MUT-7<sup>FL</sup> /MUT-8<sup>FL</sup> | 1-910 /1-578 | (N) His10-MBP-3C / (N) His6-Trx-3C | MBPTrap, 3C, Heparin, Anion exchange, SEC | Ce255 /Ce15 | Nuclease assay, Phase separation, SEC |
| MUT-7<sup>FL</sup> /MUT-8<sup>CTD</sup> | 1-910 /322-578 | (N) His10-MBP-3C / (N) His10-MBP-3C | MBPTrap, 3C, Heparin, SEC | Ce255 /Ce286 | Anisotropy |
| MUT-7<sup>D435A</sup> /MUT-8<sup>CTD</sup> | 1-910 /322-578 | (N) His10-MBP-3C / (N) His10-MBP-3C | MBPTrap, 3C, Heparin, SEC | Ce256 /Ce286 | Nuclease assay |
| MUT-7<sup>CTD</sup> /MUT-8<sup>CTD</sup> | 633-910 /322-578 | (N) His10-MBP-3C / (N) His10-MBP-3C | HISTrap, 3C, HISTrap, Anion exchange, SEC | Ce285 /Ce286 | Pulldown, Proteolysis, Crosslinking, SEC-MALS |
| MUT-7<sup>CTD(633-899)</sup> /MUT-8<sup>CTD(322-567)</sup> | 633-899 /322-567 | (N) His10-MBP-3C / (N) His10-MBP-3C | HISTrap, 3C, HISTrap, Anion exchange, SEC | Ce373 /Ce374 | Crystal screenings and structure |
| MUT-8<sup>NTD</sup> | 1-235 | (N) His10-MBP-3C | HISTrap, 3C, Anion exchange, SEC | Ce339 | CD, SEC |
| MUT-16<sup>584-724</sup> /MUT-8<sup>NTD</sup> | 584-724 /1-235 | (N) His6-GST-3C / (N) His10-MBP-3C | HISTrap, 3C, HISTrap, SEC | Ce348 /Ce339 | Phase separation, CD, SEC |
| MUT-16<sup>1-724</sup> | 1-724 | (N) GST-3C, (C) His10 | HISTrap, Anion exchange, SEC | Ce221 | Pulldown |
| MUT-16<sup>1-584</sup> | 1-584 | (N) GST-3C, (C) His10 | HISTrap, Anion exchange, SEC | Ce220 | Pulldown |
| MUT-16<sup>1-383</sup> | 1-383 | (N) GST-3C, (C) His10 | HISTrap, Anion exchange, SEC | Ce219 | Pulldown |
| MUT-16<sup>584-724</sup> | 584-724 | (N) GST-3C, (C) His10 | HISTrap, 3C, Cation exchange | Ce298 | Phase separation, CD |
| MUT-16<sup>584-724</sup> | 584-724 | (N) GST-3C, (C) His10 | HISTrap, Anion exchange, SEC | Ce298 | Pulldown |
| EXD-3<sup>FL</sup> | 1-876 | (N) His10-MBP-3C | HISTrap, Heparin, 3C, HISTrap, SEC | Hs87 | Nuclease assay |
| EXD-3<sup>NTD-EXO</sup> | 1-582 | (N) His10-MBP-3C | HISTrap, 3C, HISTrap, Heparin, | Hs88 | Nuclease assay |
|  |  |  | SEC |  |  |
| EXD-3<sup>CTD</sup> | 624-876 | (N) His10-MBP-3C | HISTrap, 3C,<br>Heparin, SEC | Hs89 | Anisotropy,<br>nuclease assay,<br>SEC-ICP-MS |
| EXD-3<sup>FL(D399A)</sup> | 1-876 | (N) His10-MBP-3C | HISTrap, 3C,<br>Heparin, SEC | Hs90 | Anisotropy |
| EXD-3<sup>NTD-EXO(D399A)</sup> | 1-582 | (N) His10-MBP-3C | HISTrap, 3C,<br>Heparin, SEC | Hs91 | Anisotropy |

**Table S3:** Constructs used for co-expression pulldown assays. Constructs only used for co-expression pulldown assays and not for protein purification are highlighted in bold. (N) and (C) indicate that the tag is N-terminally or C-terminally fused to the protein of interest, respectively. His10 or His6 indicate the presence of 10 or 6 Histidine residues, respectively. MBP= Maltose Binding Protein, Trx= Thioredoxin, GST= Glutathione S-transferase, 3C= 3C cleavage site (LEVLFQ/GP).

| Name | Residues | Tags | Plasmid # |
| --- | --- | --- | --- |
| MUT-7<sup>FL</sup> | 1-910 | (N) His10-MBP-3C | Ce255 |
| MUT-7<sup>NTD-EXO</sup> | 1-625 | (N) His10-MBP-3C | Ce333 |
| MUT-7<sup>CTD</sup> | 633-910 | (N) His10-MBP-3C | Ce285 |
| **MUT-7<sup>CTD</sup>(R853E, T855E)** | **633-910** | **(N) His10-MBP-3C** | **Ce302** |
| **MUT-7<sup>CTD-C</sup>(773-910)** | **773-910** | **(N) His10-MBP-3C** | **Ce349** |
| MUT-8<sup>FL</sup> | 1-578 | (N) His6-Trx-3C | Ce15 |
| **MUT-8<sup>FL</sup>** | **1-578** | **(N) His10-MBP-3C** | **Ce338** |
| MUT-8<sup>NTD</sup> | 1-235 | (N) His10-MBP-3C | Ce339 |
| **MUT-8<sup>CTD</sup>** | **322-578** | **(N) His10-3C** | **Ce299** |
| MUT-16<sup>1-724</sup> | 1-724 | (N) GST-3C, (C) His10 | Ce221 |
| MUT-16<sup>1-584</sup> | 1-584 | (N) GST-3C, (C) His10 | Ce220 |
| MUT-16<sup>1-383</sup> | 1-383 | (N) GST-3C, (C) His10 | Ce219 |
| MUT-16<sup>584-724</sup> | 584-724 | (N) GST-3C, (C) His10 | Ce298 |
| HsEXD-3<sup>CTD</sup> | 624-876 | (N) His10-MBP-3C | Hs89 |
| **DrEXD-3<sup>CTD</sup>** | **631-861** | **(N) His10-MBP-3C** | **Dr05** |

## Supplementary Material and Methods

### Crosslinking-Mass Spectrometry

1 mg DSBU (Disuccinimidyl Dibutyric Urea) (ThermoScientific) was resuspended in DMSO to reach a final concentration of 50 mM and stock aliquots were stored at −20°C. Increasing quantities of DSBU were added to the MUT-7^CTD^/MUT-8^CTD^ complex (9 µM) in 20 mM HEPES/NaOH pH 7.5, 150 mM NaCl, 2 mM DTT. Reactions were incubated for 30 minutes at room temperature and then stopped by adding 25 mM Tris/HCl (pH 7.5) for 15 min at room temperature. Samples were analysed by SDS-PAGE and the crosslinked band of interest was excised from the gel. The Coomassie-stained gel band was destained with a mixture of acetonitrile and 50 mM ammonium bicarbonate. The proteins were reduced using 10 mM DTT and alkylated with 50 mM iodoacetamide. Trypsin (Mass Spectrometry Grade) was used for proteolytic cleavage. Digestion was carried out with trypsin at 37°C overnight. Formic acid was used to stop the digestion and extracted peptides were desalted using C18 Stagetips_66._

Peptides were separated on a ultimate 3000 nano-flow chromatography system (Thermo-Fisher), using a pre-column for sample loading (Acclaim PepMap C18, 2 cm × 0.1 mm, 5 μm, Thermo-Fisher), and a C18 analytical column (Acclaim PepMap C18, 50 cm × 0.75 mm, 2 μm, Thermo-Fisher), applying a segmented linear gradient from 2% to 35% and finally 80% solvent B (80 % acetonitrile, 0.1 % formic acid; solvent A 0.1 % formic acid) at a flow rate of 230 nL/min over 120 minutes.

Eluting peptides were analysed on an Exploris 480 Orbitrap mass spectrometer (Thermo Fisher) using coated emitter tips (PepSep, MSWil) with the following settings: The mass spectrometer was operated in DDA mode with the FAIMS CV set to -40, -55 and -70. The cycle time was set to 1 s. The survey scans were obtained in a mass range of 375-1600 m/z, at a resolution of 120k at 200 m/z, and a normalized AGC target at 100%. The selected ions were isolated with a width of 1.2 m/z, fragmented in the HCD cell at 27%, 30%, 33% collision energy, and the spectra recorded for max. 200 ms at a normalized AGC target of 200% and a resolution of 30k. Peptides with a charge of +3 to +8 were included for fragmentation, the peptide match feature was set to preferred, the exclude isotope feature was enabled, and selected precursors were dynamically excluded from repeated sampling for 30 seconds.

Raw data were processed using the MaxQuant software package 2.0.3.0 ^6^^7^ and searched against the target sequence, the Uniprot *E. coli* reference proteome (version 2021_03, www.uniprot.org) as well as a database of most common contaminants. The search was performed with standard identification settings: full trypsin specificity allowing a maximum of two missed cleavages. Carbamidomethylation of cysteine residues was set as fixed, oxidation of methionine and acetylation of protein N-termini as variable modifications. All other settings were left at default. Results were filtered at a false discovery rate of 1% at protein and peptide spectrum match level.

To identify cross-linked peptides, the spectra were searched using Merox software 2.0 _68_ against the sequences of the top 10 non-contaminant proteins from the MQ search sorted by iBAQ. Carbamidomethylation of cysteine was set as fixed, oxidation of methionine and acetylation of protein N-termini as variable modifications. The enzyme specificity was set to trypsin allowing 4 missed cleavage sites. Crosslinker settings were selected as DSBU. Search results were filtered for 1% FDR (false discovery rate) on the PSM level (peptide-spectrum matches) and a maximum precursor mass deviation of 5 ppm. To remove low quality PSMs, additionally an score cutoff of 50 was applied. Cross-link maps were generated in xiNET ^69^.

### Limited proteolysis

The MUT-7^CTD^/MUT-8^CTD^ complex at 0.7 mg/mL was incubated with increasing concentrations of Trypsin, Elastase, Chymotrypsin and Glu C (0.02, 0.002 and 0.0002 mg/mL) in 13 µL buffer (20 mM HEPES/NaOH, 50 mM NaCl, 2 mM MgCl_2_). Reactions were incubated for 30 minutes on ice. A time course was performed with Elastase at 0.02 mg/mL. Reaction products were visualized by SDS-PAGE. The Elastase reaction products were analysed by Size Exclusion Chromatography.

### Size-exclusion chromatography coupled to multi-angle light scattering (SEC-MALS)

The molecular mass and the oligomeric state of the MUT-7^CTD^/MUT-8^CTD^ complex were determined by size exclusion chromatography (SEC) coupled to multi-angle light scattering (MALS). The complex was analysed at a concentration of 5.4 mg/mL in a buffer consisting of 20 mM Tris/HCl pH 7.5, 150 mM NaCl, and 2 mM DTT. A Superdex 75 Increase 10/300 column (GE Healthcare Life Sciences) was connected to a 1260 Infinity HPLC system (Agilent Technologies) coupled to a MiniDawn Treos detector (Wyatt Technologies) with a laser emitting at 690 nm. An RI-101 detector (Shodex) was used for refractive index measurement. Data were analysed using the Astra 7 software package (Wyatt Technologies).

### Isothermal titration calorimetry (ITC)

ITC was carried out by using a MicroCal PEAQ-ITC calorimeter (Malvern Panalytical). All samples were dialyzed against a buffer containing 25 mM Tris/HCl pH 7.5, 150 mM NaCl, 0.25 mM TCEP. The cell was filled with 46 μM MUT-8^NTD^, the syringe with 600 μM MUT-16^584–7^^24^. Titrations were carried out at 20 °C with 19 injections of 2 μL. As control, the injectant was titrated into buffer. All data were processed and curves fitted using the MicroCal PEAQ ITC software.

### Phylogenetic analysis

Orthodb v10.1 and EggNOG v5 were used to derive predicted Nibbler/Mut7 orthologs at various taxonomic levels, Interproscan v5.42-78.0 for the identification of Pfam domains within these sequences, and iTol v6 for visualization of the phylogenetic distribution of the orthologous groups on the phylogenetic species tree (PMID: 33885785).

### Size-Exclusion Chromatography-Inductively Coupled Plasma Mass Spectrometry (SEC-ICP-MS)

HsEXD3^CTD^ was purified as described in **Table S2** and stored at −70 °C in 20 mM Tris/HCl (pH7.5), 150 mM NaCl, 10% (v/v) glycerol, 0.5 mM TCEP at a concentration of 1.5 mg/mL prior to measurement. A NexSAR Speciation Analysis Ready HPLC system from Perkin Elmer (Massachusetts, USA) controlled by Clarity 8.8 software was used for the size exclusion separation. The mobile phase consisted of 75 mmol/L NaCl and 10 mmol/L Tris/HCl at pH 7.2 and the flow rate was set to 0.3 mL min^-1^. The injection volume for injection of samples, blank solutions and standards onto the ACQUITY UPLC Protein BEH 200Å SEC column (Waters) was 15 µL. For elemental detection, the system was combined with a NexION 2000 quadrupole ICP-MS from Perkin Elmer (Massachusetts, USA). The reaction cell was filled with oxygen to induce the formation of ^32^S^16^O^+^ and circumvent the ^16^O_2+_ interference on the most abundant sulfur isotope ^32^S. Zinc was detected as ^66^Zn. Bovine Cu/Zn-superoxide dismutase (SOD) was used to determine the intensity ratio between the ^66^Zn and the ^32^S^16^O signals. The stoichiometric zinc ratio was calculated via the number of sulfur atoms per protein. Details concerning the optimization of the ICP-MS parameters and data evaluation can be found in a previous publication ^70^. pH measurements were conducted with an Education Line EL20 pH-Meter with a micro electrode (inLab Micro, Mettler, Toledo, USA).

## References

1. Buchon, N. & Vaury, C. RNAi: a defensive RNA-silencing against viruses and transposable elements. Heredity 96, 195–202 (2006).

2. Gebert, D. & Rosenkranz, D. RNA-based regulation of transposon expression. WIREs RNA 6, 687–708 (2015).

3. Meister, G. Argonaute proteins: functional insights and emerging roles. Nat. Rev. Genet. 14, 447–459 (2013).

4. Ozata, D. M., Gainetdinov, I., Zoch, A., O’Carroll, D. & Zamore, P. D. PIWI-interacting RNAs: small RNAs with big functions. Nat. Rev. Genet. 20, 89–108 (2019).

5. Czech, B. et al. piRNA-Guided Genome Defense: From Biogenesis to Silencing. Annu. Rev. Genet. 52, 131–157 (2018).

6. Czech, B. & Hannon, G. J. One Loop to Rule Them All: The Ping-Pong Cycle and piRNA-Guided Silencing. Trends Biochem. Sci. 41, 324–337 (2016).

7. Feltzin, V. L. et al. The exonuclease Nibbler regulates age-associated traits and modulates pi RNA length in *D rosophila*. Aging Cell 14, 443–452 (2015).

8. Hayashi, R. et al. Genetic and mechanistic diversity of piRNA 3′-end formation. Nature 539, 588–592 (2016).

9. Wang, H. et al. Antagonistic roles between Nibbler and Hen1 modulate piRNA 3’ ends in *Drosophila*. Development dev.128116 (2015) doi:10.1242/dev.128116.

10. Almeida, M. V., Andrade-Navarro, M. A. & Ketting, R. F. Function and Evolution of Nematode RNAi Pathways. Non-Coding RNA 5, (2019).

11. Phillips, C. M., Montgomery, T. A., Breen, P. C. & Ruvkun, G. MUT-16 promotes formation of perinuclear Mutator foci required for RNA silencing in the C. elegans germline. Genes Dev. 26, 1433–1444 (2012).

12. Uebel, C. J., Agbede, D., Wallis, D. C. & Phillips, C. M. Mutator Foci Are Regulated by Developmental Stage, RNA, and the Germline Cell Cycle in Caenorhabditis elegans. G3 Bethesda Md 10, 3719–3728 (2020).

13. Uebel, C. J. et al. Distinct regions of the intrinsically disordered protein MUT-16 mediate assembly of a small RNA amplification complex and promote phase separation of Mutator foci. PLOS Genet. 14, e1007542 (2018).

14. Tsai, H.-Y. et al. A Ribonuclease Coordinates siRNA Amplification and mRNA Cleavage during RNAi. Cell 160, 407–419 (2015).

15. Preston, M. A. et al. Unbiased screen of RNA tailing activities reveals a poly(UG) polymerase. Nat. Methods 16, 437–445 (2019).

16. Chen, C.-C. G. et al. A member of the polymerase beta nucleotidyltransferase superfamily is required for RNA interference in C. elegans. Curr. Biol. CB 15, 378–383 (2005).

17. Shukla, A. et al. poly(UG)-tailed RNAs in genome protection and epigenetic inheritance. Nature 582, 283–288 (2020).

18. Roschdi, S. et al. An atypical RNA quadruplex marks RNAs as vectors for gene silencing. Nat. Struct. Mol. Biol. 29, 1113–1121 (2022).

19. Gu, W. et al. Distinct Argonaute-Mediated 22G-RNA Pathways Direct Genome Surveillance in the C. elegans Germline. Mol. Cell 36, 231–244 (2009).

20. Aoki, K., Moriguchi, H., Yoshioka, T., Okawa, K. & Tabara, H. In vitro analyses of the production and activity of secondary small interfering RNAs in C. elegans. EMBO J. 26, 5007–5019 (2007).

21. Tabara, H. et al. The rde-1 Gene, RNA Interference, and Transposon Silencing in C. elegans. Cell 99, 123–132 (1999).

22. Ketting, R. F., Haverkamp, T. H. A., van Luenen, H. G. A. M. & Plasterk, R. H. A. mut-7 of C. elegans, Required for Transposon Silencing and RNA Interference, Is a Homolog of Werner Syndrome Helicase and RNaseD. Cell 99, 133–141 (1999).

23. Zhang, C. et al. mut-16 and other mutator class genes modulate 22G and 26G siRNA pathways in Caenorhabditis elegans. Proc. Natl. Acad. Sci. U. S. A. 108, 1201–1208 (2011).

24. Tops, B. B. J. et al. RDE-2 interacts with MUT-7 to mediate RNA interference in Caenorhabditis elegans. Nucleic Acids Res. 33, 347–355 (2005).

25. Liu, N. et al. The Exoribonuclease Nibbler Controls 3′ End Processing of MicroRNAs in Drosophila. Curr. Biol. 21, 1888–1893 (2011).

26. Han, B. W., Hung, J.-H., Weng, Z., Zamore, P. D. & Ameres, S. L. The 3′-to-5′ Exoribonuclease Nibbler Shapes the 3′ Ends of MicroRNAs Bound to Drosophila Argonaute1. Curr. Biol. 21, 1878–1887 (2011).

27. Xie, W. et al. Structure-function analysis of microRNA 3′-end trimming by Nibbler. Proc. Natl. Acad. Sci. 117, 30370–30379 (2020).

28. Matelska, D., Steczkiewicz, K. & Ginalski, K. Comprehensive classification of the PIN domain-like superfamily. Nucleic Acids Res. 45, 6995–7020 (2017).

29. Iyer, L. M., Burroughs, A. M. & Aravind, L. The prokaryotic antecedents of the ubiquitin-signaling system and the early evolution of ubiquitin-like β-grasp domains. Genome Biol. 7, R60 (2006).

30. van Kempen, M. et al. Fast and accurate protein structure search with Foldseek. Nat. Biotechnol. 1–4 (2023) doi:10.1038/s41587-023-01773-0.

31. Holm, L., Laiho, A., Törönen, P. & Salgado, M. DALI shines a light on remote homologs: One hundred discoveries. Protein Sci. 32, e4519 (2023).

32. Bank, R. P. D. RCSB PDB -5TQJ: Crystal structure of response regulator receiver protein from Burkholderia phymatum. https://www.rcsb.org/structure/5TQJ.

33. Oerum, S. et al. Structural studies of RNase M5 reveal two-metal-ion supported two-step dsRNA cleavage for 5S rRNA maturation. RNA Biol. 18, 1996–2006 (2021).

34. Zhang, Y. & Skolnick, J. TM-align: a protein structure alignment algorithm based on the TM-score. Nucleic Acids Res. 33, 2302–2309 (2005).

35. Manage, K. I. et al. A tudor domain protein, SIMR-1, promotes siRNA production at piRNA-targeted mRNAs in C. elegans. eLife 9, e56731 (2020).

36. Woody, R. W. Circular Dichroism Spectrum of Peptides in the Poly(Pro)II Conformation. ACS Publications (2009) doi:10.1021/ja901218m.

37. Aoki, S. T. et al. C. elegans germ granules require both assembly and localized regulators for mRNA repression. Nat. Commun. 12, 996 (2021).

38. Shukla, A., Perales, R. & Kennedy, S. piRNAs coordinate poly(UG) tailing to prevent aberrant and perpetual gene silencing. Curr. Biol. CB 31, 4473–4485.e3 (2021).

39. Jansson-Fritzberg, L. I. et al. DNMT1 inhibition by pUG-fold quadruplex RNA. RNA N. Y. N 29, 346–360 (2023).

40. Hsu, T.-Y., Zhang, B., L’Etoile, N. D. & Juang, B.-T. C. elegans orthologs MUT-7/CeWRN-1 of Werner syndrome protein regulate neuronal plasticity. eLife 10, e62449 (2021).

41. Mori, G. et al. Rv0579 Is Involved in the Resistance to the TP053 Antitubercular Prodrug. Front. Microbiol. 11, (2020).

42. Kabsch, W. XDS. *Acta Crystallogr. D Biol. Crystallogr.* 66, 125–132 (2010).

43. Evans, P. R. An introduction to data reduction: space-group determination, scaling and intensity statistics. Acta Crystallogr. D Biol. Crystallogr. 67, 282–292 (2011).

44. Evans, P. R. & Murshudov, G. N. How good are my data and what is the resolution? Acta Crystallogr. D Biol. Crystallogr. 69, 1204–1214 (2013).

45. McCoy, A. J. et al. Phaser crystallographic software. *J. Appl. Crystallogr.* 40, 658–674 (2007).

46. Liebschner, D. et al. Macromolecular structure determination using X-rays, neutrons and electrons: recent developments in Phenix. Acta Crystallogr. Sect. Struct. Biol. 75, 861– 877 (2019).

47. Bond, P. S. & Cowtan, K. D. ModelCraft: an advanced automated model-building pipeline using Buccaneer. Acta Crystallogr. Sect. Struct. Biol. 78, 1090–1098 (2022).

48. Cowtan, K., Metcalfe, S. & Bond, P. Shift-field refinement of macromolecular atomic models. Acta Crystallogr. Sect. Struct. Biol. 76, 1192–1200 (2020).

49. Potterton, L. et al. CCP4i2: the new graphical user interface to the CCP4 program suite. Acta Crystallogr. Sect. Struct. Biol. 74, 68–84 (2018).

50. Emsley, P., Lohkamp, B., Scott, W. G. & Cowtan, K. Features and development of Coot. Acta Crystallogr. D Biol. Crystallogr. 66, 486–501 (2010).

51. Afonine, P. V. et al. Towards automated crystallographic structure refinement with phenix.refine. Acta Crystallogr. D Biol. Crystallogr. 68, 352–367 (2012).

52. Kovalevskiy, O., Nicholls, R. A., Long, F., Carlon, A. & Murshudov, G. N. Overview of refinement procedures within REFMAC5: utilizing data from different sources. Acta Crystallogr. Sect. Struct. Biol. 74, 215–227 (2018).

53. Williams, C. J. et al. MolProbity: More and better reference data for improved all-atom structure validation. Protein Sci. Publ. Protein Soc. 27, 293–315 (2018).

54. Joosten, R. P., Joosten, K., Murshudov, G. N. & Perrakis, A. PDB_REDO: constructive validation, more than just looking for errors. Acta Crystallogr. D Biol. Crystallogr. 68, 484– 496 (2012).

55. Schindelin, J., et al. Fiji: an open-source platform for biological-image analysis. *Nat. Methods* 9, 676–682 (2012).

56. Madeira, F. et al. Search and sequence analysis tools services from EMBL-EBI in 2022. Nucleic Acids Res. 50, W276–W279 (2022).

57. Waterhouse, A. M., Procter, J. B., Martin, D. M. A., Clamp, M. & Barton, G. J. Jalview Version 2—a multiple sequence alignment editor and analysis workbench. *Bioinformatics* 25, 1189–1191 (2009).

58. Brenner, S. The genetics of Caenorhabditis elegans. Genetics 77, 71–94 (1974).

59. Paix, A., Folkmann, A. & Seydoux, G. Precision genome editing using CRISPR-Cas9 and linear repair templates in C. elegans. Methods San Diego Calif 121–122, 86–93 (2017).

## Supplementary references

60. Bank, R. P. D. RCSB PDB -5TQJ: Crystal structure of response regulator receiver protein from Burkholderia phymatum. https://www.rcsb.org/structure/5TQJ.

61. Oerum, S. et al. Structural studies of RNase M5 reveal two-metal-ion supported two-step dsRNA cleavage for 5S rRNA maturation. RNA Biol. 18, 1996–2006 (2021).

62. Dar, A. C., Dever, T. E. & Sicheri, F. Higher-Order Substrate Recognition of eIF2α by the RNA-Dependent Protein Kinase PKR. Cell 122, 887–900 (2005).

63. Hollmann, N. M. et al. Upstream of N-Ras C-terminal cold shock domains mediate poly(A) specificity in a novel RNA recognition mode and bind poly(A) binding protein. Nucleic Acids Res. 51, 1895–1913 (2023).

64. Singh, S. P., Kukshal, V., De Bona, P., Antony, E. & Galletto, R. The mitochondrial single-stranded DNA binding protein from S. cerevisiae, Rim1, does not form stable homo-tetramers and binds DNA as a dimer of dimers. *Nucleic Acids Res.* **46**, 7193–7205 (2018).

65. Wu, M. et al. Structures of a key interaction protein from the Trypanosoma brucei editosome in complex with single domain antibodies. J. Struct. Biol. 174, 124–136 (2011).

66. Rappsilber, J., Mann, M. & Ishihama, Y. Protocol for micro-purification, enrichment, pre-fractionation and storage of peptides for proteomics using StageTips. Nat. Protoc. 2, 1896–1906 (2007).

67. Tyanova, S., Temu, T. & Cox, J. The MaxQuant computational platform for mass spectrometry-based shotgun proteomics. Nat. Protoc. 11, 2301–2319 (2016).

68. Iacobucci, C. et al. A cross-linking/mass spectrometry workflow based on MS-cleavable cross-linkers and the MeroX software for studying protein structures and protein-protein interactions. Nat. Protoc. 13, 2864–2889 (2018).

69. Combe, C. W., Fischer, L. & Rappsilber, J. xiNET: cross-link network maps with residue resolution. Mol. Cell. Proteomics MCP 14, 1137–1147 (2015).

70. Hann, S., Koellensperger, G., Obinger, C., Furtmüller, P. G. & Stingeder, G. SEC-ICP-DRCMS and SEC-ICP-SFMS for determination of metal–sulfur ratios in metalloproteins. J. Anal. At. Spectrom. 19, 74–79 (2004).

